# Chromatin remodelling subunit SMARCB1 is implicated in dendrite development and complex brain functions

**DOI:** 10.1101/2025.10.30.685571

**Authors:** Kristina I. Lemke, Alina Filatova, Joanna Chiang, Hannah North, Myrthe R.M. Kamphof, Michaela Becker-Röck, Bodo Laube, Gijs W. E. Santen, Hanna Swaab, Ulrike A. Nuber

**Author notes:** These authors contributed equally to this work (Kristina I. Lemke, Alina Filatova, Joanna Chiang). Correspondence to: Professor Ulrike A. Nuber Full address: Stem Cell and Development Biology, Technical University of Darmstadt, Schnittspahnstraße 13, 64287 Darmstadt, Germany.

## Abstract

*SMARCB1* encodes a core component of the BAF chromatin remodelling complex and pathogenic variants in this gene are associated with neurodevelopmental disorders such as Coffin-Siris syndrome. The relationship between altered SMARCB1 protein products and severe functional brain changes in Coffin-Siris syndrome remains largely unknown. We performed cellular, molecular, and behavioural analyses of a Coffin-Siris syndrome mouse model with a heterozygous nervous system-specific *Smarcb1* mutation. In addition, we evaluated general cognitive abilities, as well as cognitive and behavioural functioning, in individuals with SMARCB1-related Coffin-Siris syndrome. *Smarcb1* mutant mice exhibited deficits in fine motor coordination and balance, as well as impaired spatial learning and memory. Furthermore, these mice showed anxiety-like behaviours and agitation when exposed to novel environments. The detected behavioural abnormalities could indicate impaired decision-making, which results in impaired risk assessment. Comparable cognitive and behavioural deviations were identified in individuals with Coffin-Siris syndrome and *SMARCB1* pathogenic variants. Our analysis of the *Smarcb1* mouse model revealed structural alterations in the brain, including decreased dendritic length and complexity of dendritic trees. These alterations may explain the observed functional impairments. Notably, our finding of reduced *Wasl* transcripts in mutant Purkinje cell nuclei suggests that dysregulation of actin polymerization may be involved in the discovered dendritic defects. Taken together, we demonstrate a link between the chromatin remodelling complex component SMARCB1, complex brain functions, neuronal structure, and a key regulator of actin branching.

## Introduction

Pathogenic variants in genes encoding components of multiprotein BAF (BRG1/BRM-associated factor; mammalian SWI/SNF) ATP-dependent chromatin remodelling complexes cause a variety of neurodevelopmental disorders, including Coffin-Siris syndrome, Nicolaides– Baraitser syndrome, and autism spectrum disorders (for review, see [2, 55]), and together represent one of the most common causes of intellectual disability (ID). A hallmark of these entities, sometimes referred to as BAFopathies, is impaired central nervous system function, manifested as diverse cognitive, behavioural, and motor phenotypes including intellectual disability, hypotonia, and difficulties in social interactions. Varying degrees of gross morphological brain alterations, such as microcephaly and brain midline abnormalities are present in some of the BAF-related disorders. However, functional relationships between the mutated genes and associated phenotypes remain unclear, particularly at the cellular and neuronal network levels.

Normal brain function relies on the establishment and maintenance of accurate neuronal connections – dendrites, axons, and their synapses – for the transmission of signals. Dendrites, the main receiving neuronal processes, are typically highly branched structures and in vertebrates are predominantly formed during the early postnatal period [47]. Purkinje cells, located in the middle layer of the cerebellar cortex, and being the sole output of this brain region with their axons projecting to cerebellar nuclei, possess a unique morphology and are an attractive model system for dendrite development. They have a strikingly large soma and nucleus and long, extensively branched dendrites that form an extremely flat, almost two-dimensional tree aligned along the sagittal cerebellar axis in the outer layer of the cerebellar cortex [26, 37, 73]. This elaborate dendritic tree is optimal to form a very high number of connections with perpendicularly running axons (called parallel fibres) derived from granule cells, the cell bodies of which demarcate the inner layer of the cerebellar cortex. Dendrite morphogenesis is regulated by cell-extrinsic and -intrinsic factors, and the stereotypical emergence of dendritic architectures denotes a tightly controlled molecular program underlying this process [28, 45, 46, 61, 73]. Concerning cell-intrinsic factors, only a few transcription factors are known to control dendrite morphogenesis in vertebrates; however, a growing number of protein components of ATP-dependent chromatin remodelling complexes have been found to play a role in this process [4, 9, 17, 19, 34, 36, 80, 81, 84]. These complexes contain an ATPase subunit of the SNF2 superfamily of proteins [23], and based on the further diversification with respect to the ATPase, four subcategories of complexes are distinguished in eukaryotic cells, ISWI, CHD, INO80, and SWI/SNF. Mammalian SWI/SNF complexes are defined by the presence of one of the mutually exclusive ATPases BRG1 (current protein name: SMARCA4) and BRM (current protein name: SMARCA2), underlying their designation BAF (BRG1/BRM-associated factor) complex. Different BAF chromatin remodelling complexes exist, and according to their composition of various constitutive and accessory protein subunits, the canonical BAF (cBAF), the polybromo-associated BAF (PBAF) and the non-canonical ncBAF/GBAF type are distinguished. These types can occur in a cell-type dependent manner and are thought to determine functional differences underlying cellular development into specific lineages [2, 33, 51, 65]. SMARCB1 is a core subunit of cBAF and PBAF complexes, and binds to the acidic patch of a nucleosome via its C-terminal basic alpha-helix [3, 32, 52]. Structurally, BAF complexes can be divided into three modules, consisting of one (the ATPase module) or multiple protein components (actin-related protein (ARP) and base modules), with SMARCB1 being part of the base module [32]. Individual genes encoding components of ATP-dependent chromatin remodelling complexes have been shown to play a role in dendrite development, particularly in mouse models. These include BAF complex component genes *Arid1b* [36], *Smarca4* [17, 34], *Actl6b* [9, 81]*, Bcl7a* [80] and *Ss18l1* (*Crest)* [1]. In addition, genes encoding proteins of other ATP-dependent chromatin remodelling complex families, the ISWI family (*Smarca5,* [4]) and the CHD family of chromatin remodelling complexes (*Chd4, Chd5,* and *Chd8*) [19, 59, 83, 84] are implicated in mammalian dendrite development (Supplementary Table 1). Pathogenic variants in six of these seven genes cause neurodevelopmental disorders in humans: *ARID1B*, *SMARCA4*, *ACTL6B, SMARCA5, CHD4,* and *CHD8* [48, 55]. Although gross cognitive, behavioural, and motor phenotypes have been described in individuals with these and related neurodevelopmental disorders caused by monoallelic pathogenic variants in genes encoding chromatin remodelling complex components, detailed phenotypic information on affected persons remains limited. However, this is an essential basis for elucidating why and how human brain development is sensitive to gene alterations encoding specific components of different chromatin remodelling complex classes. Ultimately, the entire chain of causal and complex relationships from these specific disease genes, their molecular and cellular consequences to higher brain functions needs to be clarified and this knowledge will contribute to improved therapeutic interventions.

Here we employ a mouse model with a heterozygous, nervous system-specific partial loss-of-function mutation in *Smarcb1* to understand how this gene is linked to impaired brain function characteristic for *SMARCB1*-related human disorders. As previously described, these mice show features of two neurodevelopmental disorders caused by *SMARCB1* pathogenic variants in humans: Coffin-Siris-syndrome and *SMARCB1*-related intellectual disability with choroid plexus hyperplasia (ID-CPH) [18, 24, 74]. In this study, we demonstrate that *Smarcb1* is involved in dendrite development in *Smarcb1* mutant mice and we describe behavioural, cognitive, and motor abnormalities of these animals. The findings are supported by behavioural analyses in humans with *SMARCB1*-related Coffin-Siris syndrome. Our data provide new insights into the consequences of *Smarcb1* mutations on neuron development, cognition, behaviour, and motor skills, establishing a link between this gene, dendrite-based neuronal connectivity, and higher brain function.

## Materials and methods

Materials and methods are provided in the Supplementary material.

## Results

### Impaired cerebellum development in *Smarcb1* mutant mice

Mice with a heterozygous nervous system-specific partial loss-of-function mutation in *Smarcb1* (*Smarcb1^+/inv^ NesCre^+/−^*) are characterized by brain midline abnormalities, microcephaly, various cerebellar alterations, and choroid plexus hyperplasia, recapitulating findings in human individuals with SMARCB1-related Coffin-Siris syndrome and with ID-CPH [24]. These phenotypes are present in varying degrees in this mouse model and are caused by a reduction of *Smarcb1* expression. The *Cre*-mediated inversion of *Smarcb1* exon 1 takes place in *Nestin*-expressing cells, the majority being radial glial cells, i.e. neural stem cells, and results in an inactivation of one *Smarcb1* allele. The inversion is reversible and continues back and forth as long as *Cre* is expressed, theoretically leading to about 50% of the *Nestin*-expressing cells and also their progeny, neurons and glial cells, possessing an active *Smarcb1* allele and 50% possessing an inactive one, in addition to the second wild-type allele. *Smarcb1^+/inv^ NesCre^+/−^* animals are referred to as mutants and *Smarcb1^+/inv^ NesCre^−/−^* are named controls since they lack the *NesCre* transgene and do not show any obvious differences to wild-type mice [24]. In our previous study, we described gross macroscopic and histological aberrations of this mouse model. Cerebellar alterations had been analysed until postnatal week 3-4 and included cerebellar hypoplasia, reduced lobulation, vermis hypoplasia, and fusion of cerebellar hemispheres [24]. Mammalian cerebellar development is only completed in the late postnatal stage and dendritic arbors of Purkinje cells reach their maximal length by P20 in mice [73]. Previously, we found that about 80% of mutant animals died before postnatal week 9, and such animals typically showed more severe brain phenotypes [24]. In this work, we focused on animals with milder phenotypes to understand how a reduced *Smarcb1* expression in Nestin-positive cells and their neuronal progeny leads to cerebellar abnormalities.

To investigate the localization of SMARCB1 protein at different stages of cerebellum development, we performed immunostainings of cerebellar tissue sections from E16.5, P7, P10, and P21 mutant and control mice (Fig. 1a-l, t-é). In all cases, sections through the middle part of the cerebellum, the vermis, were investigated. In addition to SMARCB1, we investigated the tissue distribution of calbindin 1 (calbindin-D28k), a protein present in Purkinje cells. These cells are derived from ventricular zone progenitors, and in rats begin to produce calbindin 1 around E14 to E15 after exiting the cell cycle, remaining migratory and having not yet reached their final position between the molecular (ML) and granular cell (GCL) layers in the mature cerebellum [22]. Immunostaining analyses revealed that SMARCB1 is present in nuclei of all cerebellar cell types in all investigated stages (E16.5, P7, P10, P21) in both mutant and control animals (Fig. 1a-d, t-w). Particularly strong SMARCB1 immunosignals were detected in Purkinje cell nuclei of P21 mouse brains in comparison to nuclei of granular cells (Fig. 1s, l’). From E16.5 to P21, the cerebella of mutant animals were consistently smaller and displayed fewer lobules than those of control mice (Fig. 1m-p, f’-i’). The cerebellar hypoplasia in mutant animals persisted into later adult stages, as shown by measurements of the cerebellar section area in five-month-old animals (Fig. 2a, b). Moreover, fewer lobules were detected in these mutant animals (Fig. 2a). The total cerebellar section area and the section areas of the GCL and ML were reduced in mutants compared to controls (Fig. 2b, d). The relative proportion of GCL and ML areas in relation to the total cerebellar area did not differ (Fig. 2e). The overall reduced cerebellum size is consistent with fissure lengths between lobules being significantly shorter or showing a tendency towards a reduced length in mutant cerebella (Fig. 2c). Collectively, these results demonstrate that SMARCB1 protein is ubiquitously localized in cerebellar tissue throughout development, with high levels present in Purkinje cells, and that *Smarcb1* mutant mice display cerebellar hypoplasia from an early developmental stage on. A hypoplastic cerebellum has also been reported in conditional *Smarcb1* knockout mice which differ from the one used in this study [54].

**Fig. 1.**
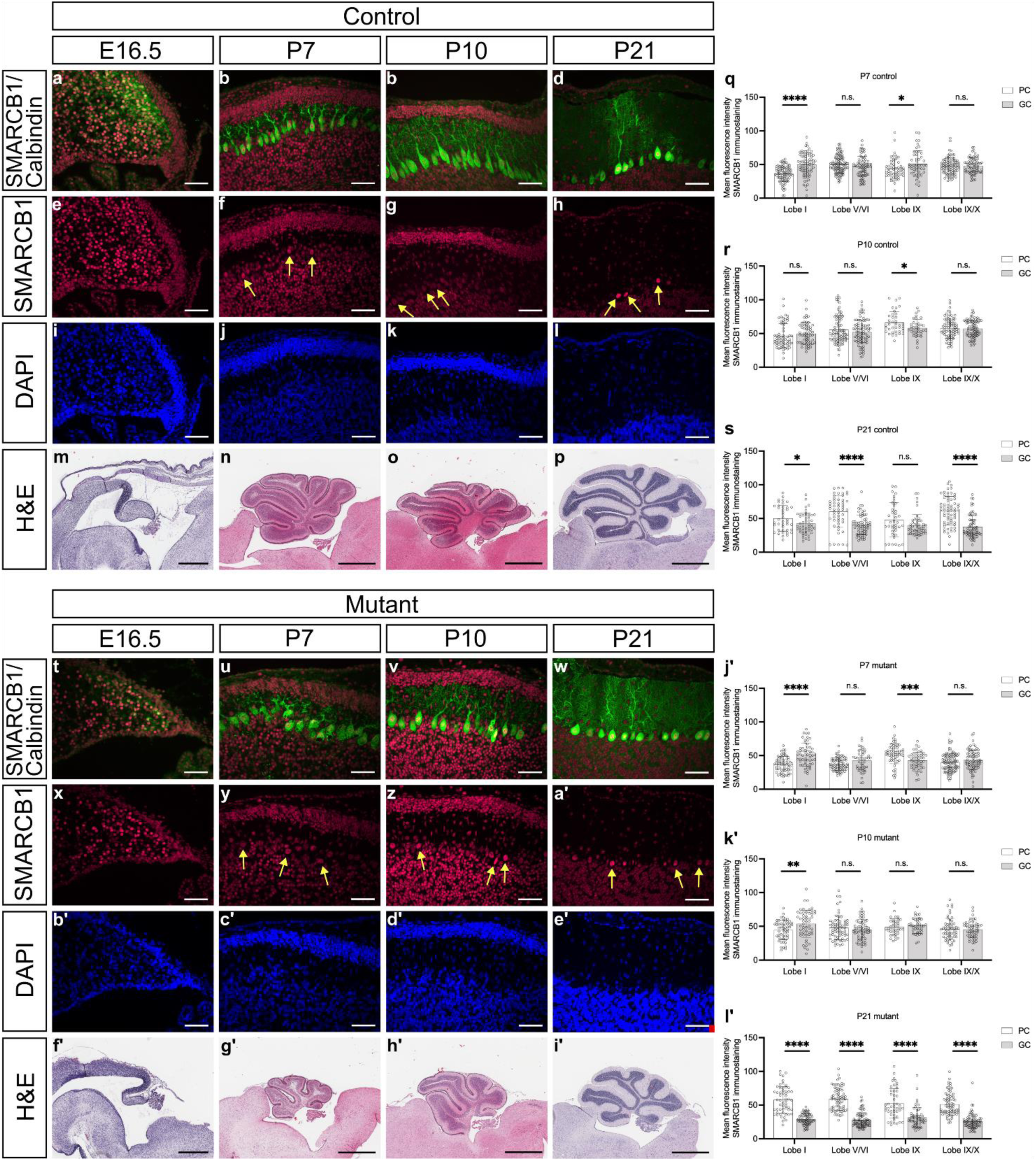
Immunofluorescence staining, SMARCB1 fluorescence intensity measurements, and hematoxylin and eosin (H&E) staining of sagittal cerebellar sections. (**a-l**, **t-e’**) Immunofluorescence and (**m-p**, **f’**-**i’**) H&E staining of sagittal vermis sections from E16.5, P7, P10, and P21 control and mutant mice. SMARCB1 (red), calbindin (green), and DAPI (blue) were stained. Yellow arrows indicate Purkinje cells. Scale bars immunofluorescence staining: 50 µm, scale bars H&E staining (P7, P10, P21): 1 mm, scale bars H&E staining (E16.5): 0.5 mm. Graphs show mean fluorescence intensity in Purkinje (PC) and granule cells (GC) in lobules I, V-VI, IX, and IX-X of (**q** and **j’**) P7, (**r** and **k’**) P10, and (**s** and **l’**) P21 control and mutant mice. Three biological replicates were used in three different experiments (three technical replicates each). Error bars mark standard deviation. * *p* ≤ 0.05, ** *p* ≤ 0.01, *** *p* ≤ 0.001, n.s. – non-significant.

**Fig. 2.**
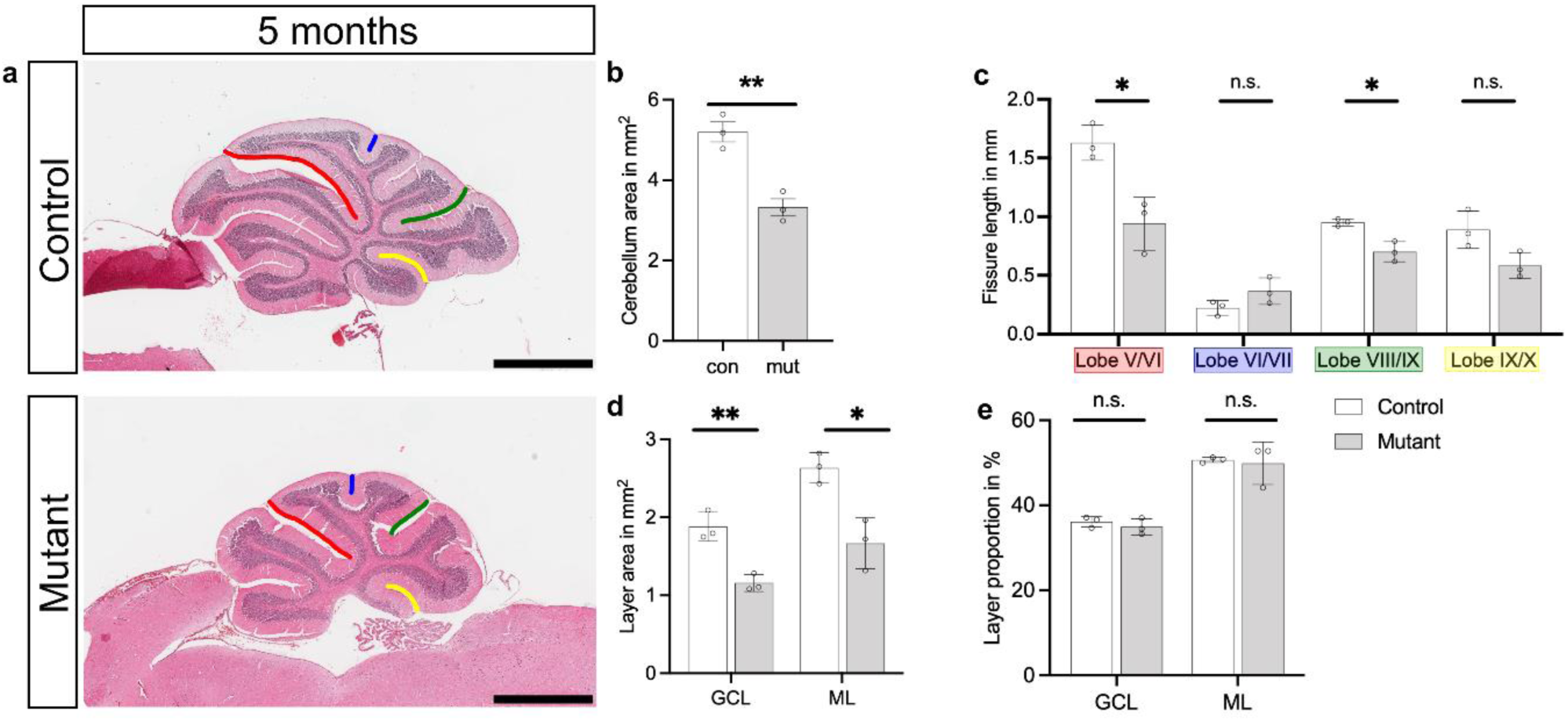
H&E staining and analysis of adult sagittal cerebellar sections. **a** H&E staining of sagittal cerebellar sections of adult (5-month-old) control and mutant mice. Scale bar: 1 mm. **b** Mean cerebellar area of control and mutant sections. **c** Graphs showing the fissure length between lobules V and VI (red), VI and VII (blue), VIII and IX (green), and IX and X (yellow). Corresponding fissures are marked in H&E staining (a). **d** Quantification of the granule cell layer (GCL) and molecular layer (ML) areas in cerebellar sections from control and mutant mice. **e** Proportion of GCL and ML areas, expressed as a percentage of the total cerebellar area, in control and mutant mice. Three biological replicates are shown. * *p* ≤ 0.05, ** *p* ≤ 0.01, *** *p* ≤ 0.001, n.s. – non-significant.

### Impaired dendrite development in *Smarcb1* mutant mice

In P7 and P10 brain sections, stages at which dendrites of individual Purkinje cells can be detected by immunostaining due to their relatively low branching density, calbindin immunosignals indicated aberrant dendritic trees in mutant animals compared to controls (Fig. 3a, b). Dendritic arborization in Purkinje cells follows a highly regulated process that undergoes distinct phases during early development. Most of the dendritic branching occurs postnatally, particularly as the neurons start to integrate into the cerebellar circuitry [7, 20, 37]. At birth (P0), mouse Purkinje cells have a simple, immature morphology with only a few short, rudimentary primary dendrites. Upon completion of migration and reaching their final position between the GCL and the ML around P4, mouse Purkinje cells form an apical pole and one to three primary stem dendrites by P8 [73]. During the transition phase, that lasts until P10, Purkinje cell dendrites start growing into the ML. Between P10 and P14, the dendritic trees mainly extend laterally, followed by a vertical expansion from P15 on. This phase is marked by an increase in both the number and length of dendritic branches. The dendritic tree becomes more elaborate, with secondary and tertiary branching now well-developed, and reaches its maximal length by P20 in mice [73]. Image analyses of paraffin brain sections stained with anti-calbindin antibodies revealed an aberrant dendrite development already at P7 (Fig. 3a). The Purkinje cell dendrites of mutant animals had a significantly shorter maximal length and the number of branches at 60 µm distance from the Purkinje cell soma was decreased. At P10, when Purkinje cells from both mutant and control animals showed a single primary stem dendrite, the dendrites of Purkinje cells of mutant animals were significantly shorter, the number of terminal dendritic ends was reduced, and primary dendrites were narrower in comparison with the control animals (Fig. 3b).

**Fig. 3.**
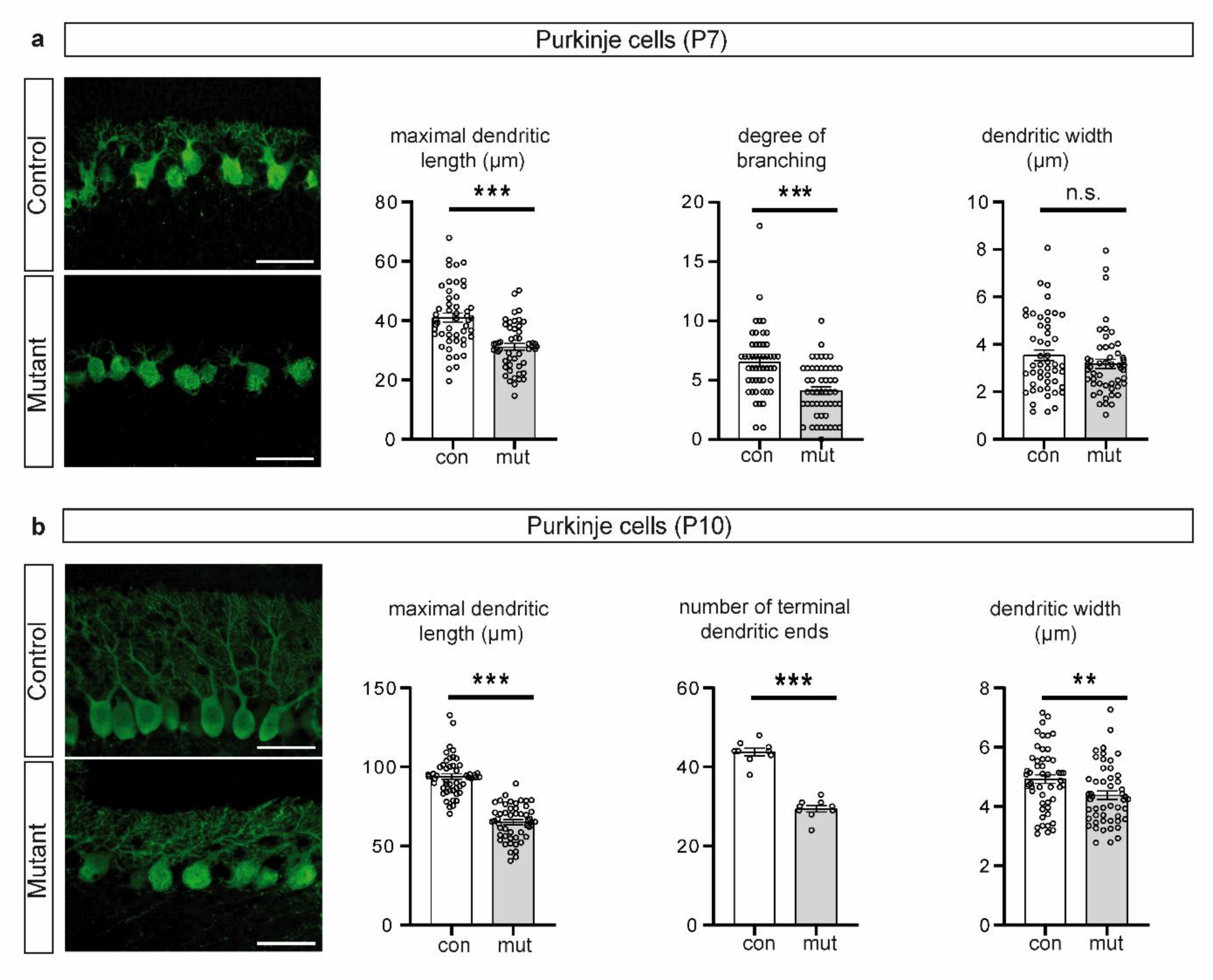
Calbindin immunosignals and dendritic morphology of Purkinje cells in (a) P7 and (b) P10 control and mutant mice. **a** Representative images of Purkinje cell immunofluorescence stainings in cerebella of P7 control and mutant mice using calbindin antibodies. Maximal dendritic length, the degree of branching (number of dendrites intersecting with a circle with diameter of 60 µm placed on the centre of the soma), and width of primary dendrites 4 µm apart from their soma origin in Purkinje cells of P7 mutant (n=3) and control (n=3) mice (50 cells each). **b** Representative images of Purkinje cell immunofluorescence stainings in cerebella of P10 control and mutant mice using calbindin antibodies. Maximal dendritic length, the number of terminal dendritic ends, and the width of primary dendrites 8 µm apart from their soma origin in Purkinje cells of P10 control (n=3) and mutant (n=3) mice (50 cells each). Scale bars, 50 µm. Error bars indicate standard error of the mean. * *p* ≤ 0.05, ** *p* ≤ 0.01, *** *p* ≤ 0.001, n.s. – non-significant.

The differences observed at early postnatal stages translated into corresponding differences in adult animals. Golgi-Cox staining of cryosections from 32-weeks-old adult mice revealed an abnormal dendrite morphology in Purkinje cells of mutant animals: dendritic length and width, the number of terminal ends, and the degree of branching were significantly reduced in comparison with controls (Fig. 4a). In summary, our data show that Purkinje cells in mutant mice have an aberrant dendrite arborization, which is already evident at P7 and persists in adults. To understand whether the aberrations in dendritic morphology observed in Purkinje cells occur in other neurons of the mutant animals and represent a general phenomenon, we analysed the dendritic morphology of neurons in the hippocampus and cerebral cortex. Pyramidal neurons in the mutant hippocampal CA1 region had a reduced number of primary basal branches and were shorter than in brain tissue from control animals (Fig. 4b). The width of primary dendrites did not differ between mutant and controls (Fig. 4b). The dendrites of granule neurons in the supra-pyramidal blade of the dentate gyrus were significantly shorter and had a significantly reduced number of branches. The width and the number of primary dendrites remained unchanged in mutant compared with control animals (Fig. 4c). Pyramidal neurons in cortical layers III and V of the mutant animals were narrower and had significantly lower arborization as evidenced by significantly lower number of primary basal branches and lower basal branching (Fig. 4d). Taken together, our data suggest that neurons in different brain regions of mutant mice have defects in dendritic morphology, decreased dendritic lengths and complexity of dendritic trees. This could impair neuronal cell and network function and, thus, lead to cognitive and behavioural deficits.

**Fig. 4.**
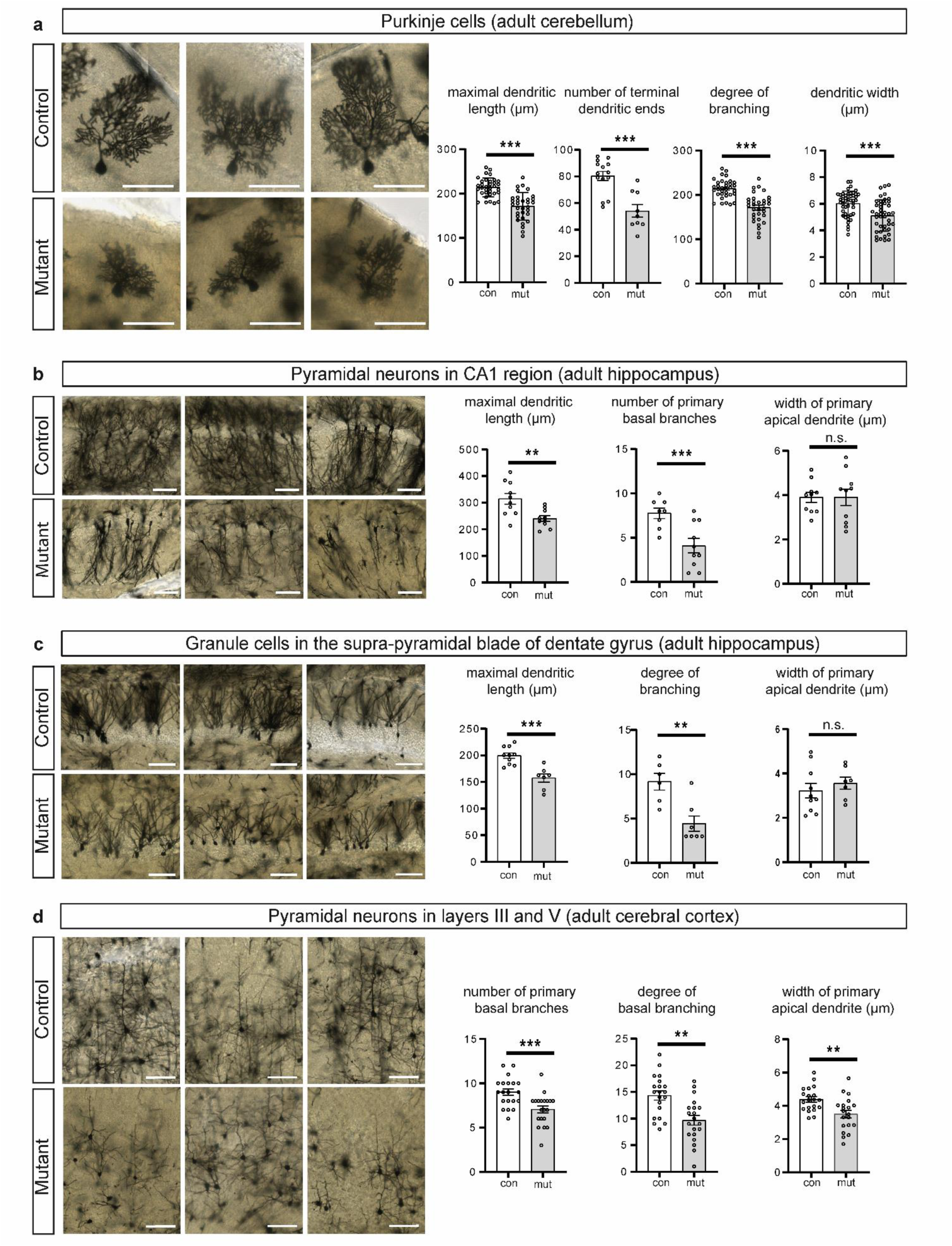
Golgi-Cox staining of neurons in different brain regions and their dendritic morphology in adult control and mutant mice. **a** Individual Purkinje cells and graphs depicting maximal dendritic length, the number of terminal dendritic ends, the degree of branching, and the width of primary dendrites 10 µm apart from their soma origin in Purkinje cells of adult control and mutant mice. n=49 (control), n=42 (mutant). **b** Pyramidal neurons in the CA1 region of the hippocampus and graphs showing maximal dendritic length, the number of primary basal branches, and the width of primary apical dendrites 10 µm apart from their soma origin in these neurons in control and mutant mice. n=10 each. **c** Granule cells in the supra-pyramidal blade of dentate gyrus and graphs depicting maximal dendritic length, the degree of branching, and the width of primary apical dendrites 10 µm apart from their soma origin in granule cells in adult control and mutant mice. n=10 (control), n=7 (mutant). **d** Pyramidal neurons in layers III and V of cerebral cortex and graphs showing the number of primary basal branches, the degree of basal branching, and the width of primary apical dendrites 10 µm apart from their soma origin in adult control and mutant mice. n=20 each. The degree of dendritic branching was determined as the number of dendrites intersecting with a circle of 80 µm diameter (pyramidal neurons in cortical layers III and V), 90 µm (Purkinje cells) or 100 µm (granule neurons located in the supra-pyramidal blade of the dentate gyrus) placed at the centre of the soma. Scale bars, 100 µm. Error bars indicate standard error of the mean. * *p* ≤ 0.05, ** *p* ≤ 0.01, *** *p* ≤ 0.001, n.s. – non-significant.

### Gene expression differences between mutant and control Purkinje cell nuclei

To gain insight into mechanisms underlying the impaired dendrite development upon partial loss-of-function mutation in *Smarcb1*, we compared candidate gene expression in Purkinje cells from control and mutant mice. Since it is very difficult to isolate whole Purkinje cells, we adapted a flow cytometry-based protocol to isolate their nuclei from cerebella of P7 animals for mRNA analysis [8]. Given the low tissue fraction of Purkinje cell nuclei, this method allowed to analyse a limited number of genes by qRT-PCR besides *Gapdh* and *Tbp* that served as reference genes. Significantly higher *Calbindin* transcript levels were determined by qRT-PCR in isolated Purkinje cell nuclei from P7 control and mutant mouse cerebella compared to all remaining nuclei, confirming the enrichment of these fractions (Fig. 5a). *Smarcb1* gene expression was lower in Purkinje cell nuclei from mutant mice than from controls, which is in line with the consequences of the *Smarcb1* mutation in these animals [24] and supports persistent effects of the genome alteration in postnatal tissue (Fig. 5b). Moreover, transcript levels of two other candidate genes were analysed: *Gap43*, a neurite outgrowth gene with reduced levels in the hippocampus of P16 *Baf53b^-/-^* mice [81], and *Wasl* that encodes the Neural Wiskott-Aldrich Syndrome Protein (N-WASP) which activates the actin nucleator ARP2/3 complex. N-WASP, together with ARP2/3, controls dendritic branching and elongation in various types of neurons, and N-WASP is crucial for Purkinje cell dendrite development *in vivo* [31]. While transcript levels of *Gap43* did not differ between mutant and control Purkinje cell nuclei, *Wasl* level were significantly reduced in the nuclei from mutant mice (Fig. 5c, d).

**Fig. 5.**
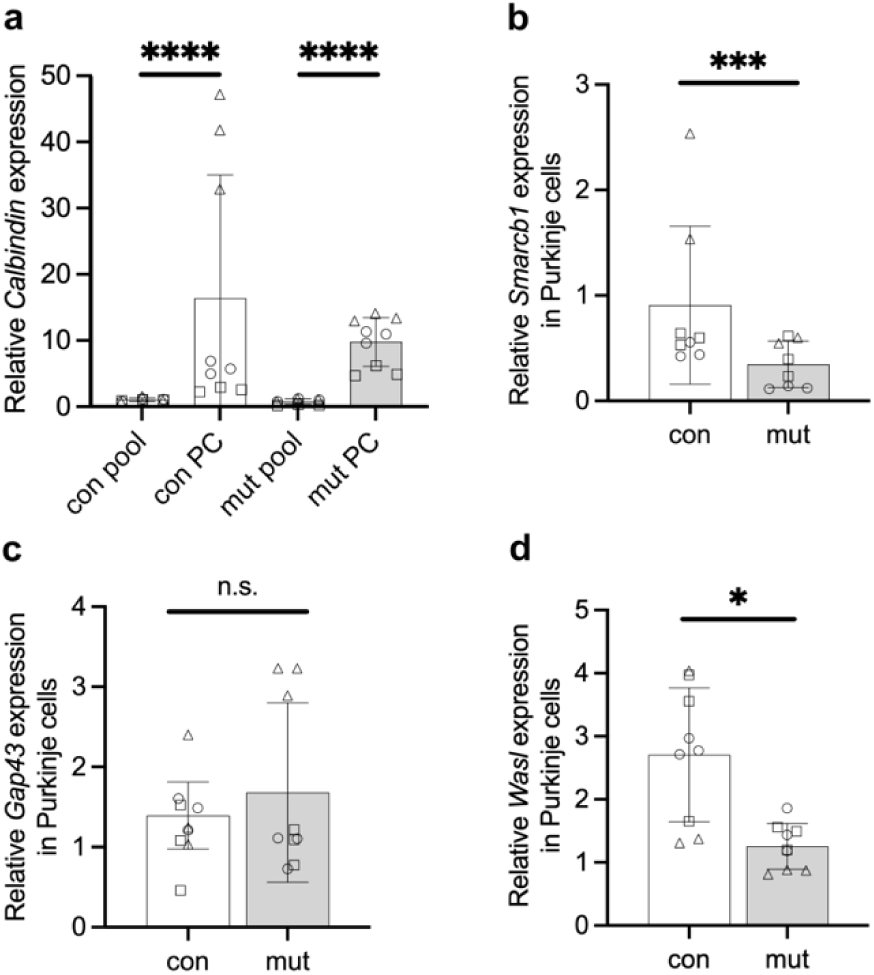
Gene expression levels in sorted Purkinje cell nuclei and remaining pooled cerebellar cells of P7 control and mutant mice, determined by RT-qPCR. **a** Gene expression levels of *Calbindin* in material from Purkinje cell (PC) and remaining pooled cerebellar cell (pool) nuclei. **b-d** Gene expression levels of *Smarcb1* (**b)**, *Wasl* (**c**), and *Gap43* (**d**) in P7 Purkinje cell nuclei. Circles, triangles, and boxes indicate cerebellum tissue from three independent litters; three technical replicates are shown each; error bars display standard deviation. * *p* ≤ 0.05, ** *p* ≤ 0.01, *** *p* ≤ 0.001, n.s. – non-significant.

### *Smarcb1* mutant mice show motor, cognitive, and behavioural abnormalities

To determine behavioural conspicuities of this mouse model, we analysed mutant and control animals that exhibited milder phenotypes, characterized by less severe brain alterations and no premature death [24]. Motor functions were assessed using the rotarod test, beam walking test, and gait analysis. Cognitive function was measured using the Morris water maze test. Anxiety-related behaviours were evaluated through the open field and elevated zero maze tests, while the visual cliff test was used to assess depth perception and visual acuity. Mutant mice did not show gross motor impairments in the rotarod test (Fig. 6a, b) and did not display any obvious gait alterations, ataxia, or tremor; their stride length was not different from control animals when considering their reduced body length (Fig. 6c, d). However, in comparison to control mice, mutant animals showed a poorer performance in the beam walking test, which measures fine motor coordination and balance (Fig. 6e-g; Supplementary Fig. 1). They took a longer time to cross the beam and showed an increased number of foot slips (Fig. 6e, f; Supplementary Fig. 1a, b). In addition, mutant mice dipped their heads more frequently during the crossing (Fig. 6g; Supplementary Fig.1c).

**Fig. 6.**
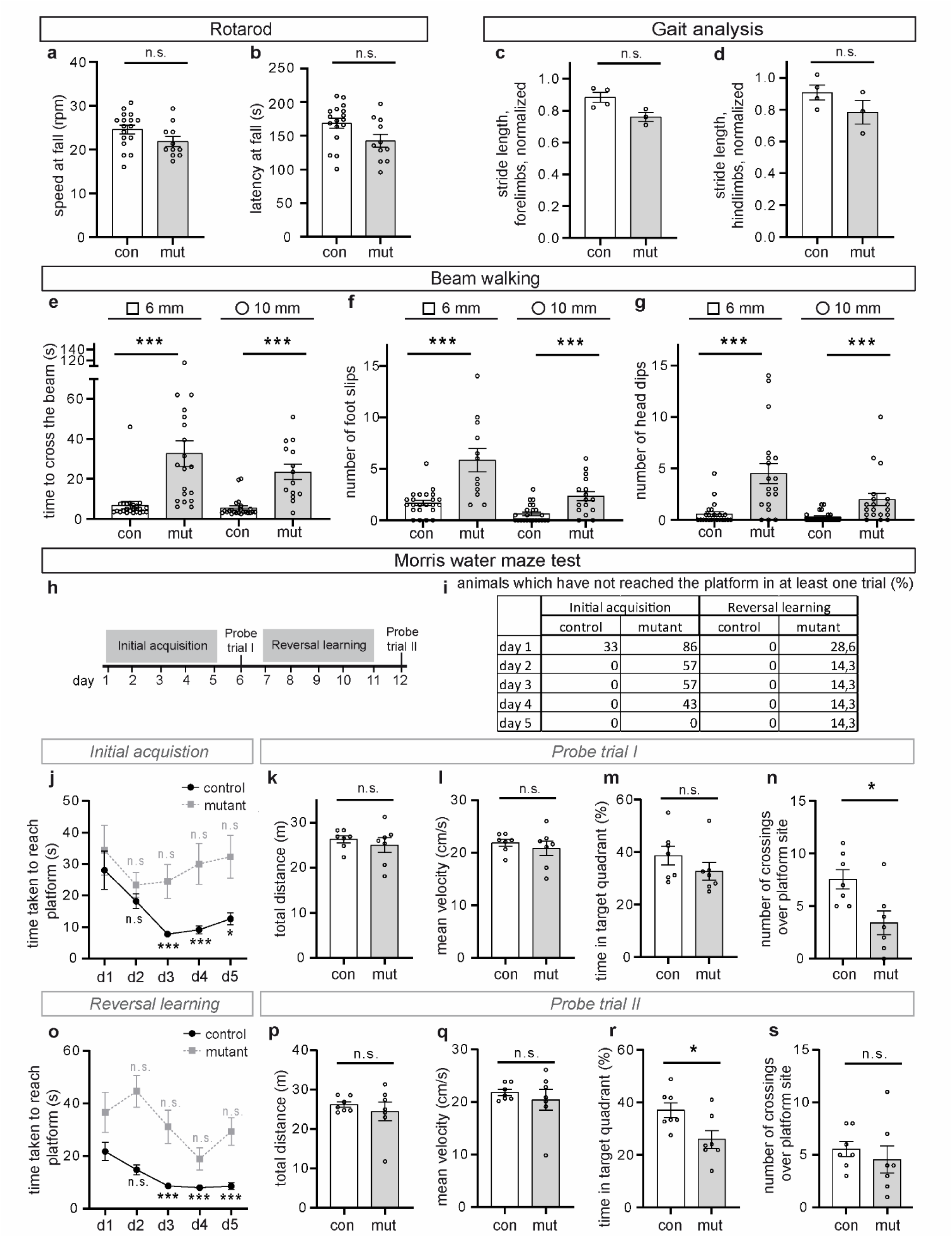
Motor function, spatial learning, and memory assessment of adult control and mutant mice **a,b** Rotarod test: speed at fall (**a**) and latency (**b**) at fall, n=17 (control), n=11 (mutant). **c**,**d** In the gait analysis, stride length was normalized to body length for forelimbs (**c**) and hindlimbs (**d**), n=4 (control), n=3 (mutant). **e-g** Beam walking test using 6 mm square and 10 mm round beams: time to cross the beam (**e**), n=23 (control), n=19 (mutant, 6 mm square beam), n=14 (mutant, 10 mm round beam); number of foot slips (**f**), n=23 (control), n=12 (mutant, 6 mm square beam), n=16 (mutant, 10 mm round beam), and head dips (**g**), n=23 (control), n=19 (mutant). **h**-**s** Morris water maze test: scheme of the experiment (**h)**, percentage of animals that did not reach the platform in at least one of four trials (**i**), n=6 (control), n=7 (mutant), time taken to reach the platform during the initial acquisition (**j**), n=28 (control), n=28 (mutant), and the reversal learning (**o**), n=28 (control), n=28 (mutant), total distance (**k**, **p**), n=7 (control), n=7 (mutant), mean velocity (**l, q**), n=7 (control), n=7 (mutant), time in target quadrant (**m, r**), n=7 (control), n=7 (mutant), and number of crossings over the platform site during probe trials I and II (**n, s**), n=7 (control), n=7 (mutant). In **j** and **o**, the time taken to reach the platform on each day was compared with that on day 1 for the respective animal group (control or mutant). Graphs in panels **k**-**n** and **p**-**s** display results from probe trials I and II from the Morris water maze test, respectively. Error bars indicate standard error of the mean. * *p* ≤ 0.05, ** *p* ≤ 0.01, *** *p* ≤ 0.001, n.s. – non-significant.

The Morris water maze test was used to assess spatial learning and memory (Fig. 6h-s). The mice underwent a series of training sessions in a water tank with a platform submerged slightly below the water surface which they had to reach by swimming. Upon platform removal, they were investigated for their ability to navigate to the former platform location using spatial cues for orientation (probe trial I). Following this initial acquisition and probe trial I, the platform was relocated to the opposite quadrant, and the time taken by the mice to find the newly positioned platform was investigated (reversal training). After 5 days, a probe trial II was conducted to determine whether the mice had learned the new location of the platform (Fig. 6h). Mutant animals did not show any gross motor impairments in swimming, as their total distance covered and mean velocity during the trials were not significantly different from control animals (Fig. 6k, l, p and q). During the initial acquisition, a significant proportion of mutant mice were not able to locate the platform within the test time of 120 seconds, even after several days of training (Fig. 6i, left columns), and in the case of successful trials, the time they took to reach the platform did not decrease, in contrast to the control mice (Fig. 6j). The poorer performance of mutant mice could be attributed to the exploration strategies that they employed to locate the platform. Whereas control animals predominantly utilized more cognitively advanced exploration strategies, namely spatial strategies that incorporate spatial cues for orientation, mutant mice more frequently used simple search strategies, such as thigmotaxis or non-spatial, i.e., random exploration strategies, even in later trials (Supplementary Fig. 2). The non-spatial exploration strategies involve simple swim paths and are associated with a low probability of locating the platform [12, 30]. In probe trial I, mutant mice spent a comparable amount of time in the target quadrant as the control mice (Fig. 6m), but they crossed the platform site significantly less often (Fig. 6n). The deficiency of mutant mice in learning the spatial cues was also apparent during the reversal learning when the platform was relocated. While control mice required less time to reach the platform during the training phase, the time required by mutant mice did not change significantly from day to day (Fig. 6o) and a proportion of mutant mice were unable to locate the platform at all within the test time of 120 seconds (Fig. 6i, right columns). As during the initial acquisition, mutant mice predominantly employed non-spatial exploration strategies (Supplementary Fig. 2). In probe trial II, mutant mice spent significantly less time in the target quadrant than control animals (Fig. 6r). Taken together, the Morris water maze test indicated that mutant mice exhibited impaired spatial learning and memory, which could be at least partially attributed to their reliance on more simple non-spatial exploration strategies.

Both the open field and elevated zero maze test results indicated an abnormal behaviour of mutant animals (Fig. 7a-d, g-k). These mice showed an increased agitation in both unfamiliar test environments, as evidenced by increased total distance covered and mean velocity (Fig. 7a, b, g and h). In the open field, mutant mice spent more time in the peripheral area and less time in the centre of the open field (Fig. 7c, d). Avoidance of the centre, i.e. a large open area, is considered as anxiety-related [75]. Conversely, in the elevated zero maze test, mutant mice spent more time in the open, unprotected arms than the control mice (Fig. 7i). While in the unprotected arms, mutant mice exhibited a higher frequency of head-dipping (downward movements of the head toward the floor) (Fig. 6j) and made extreme vertical dips, resulting in falls from the apparatus (Fig. 7k). Head dips are associated with decreased anxiety and with an exploratory/risk assessment behaviour, and anxiety-reducing medication leads to an increase in time spent in the open arms and increased frequency of head-dipping [11, 71].

**Fig. 7.**
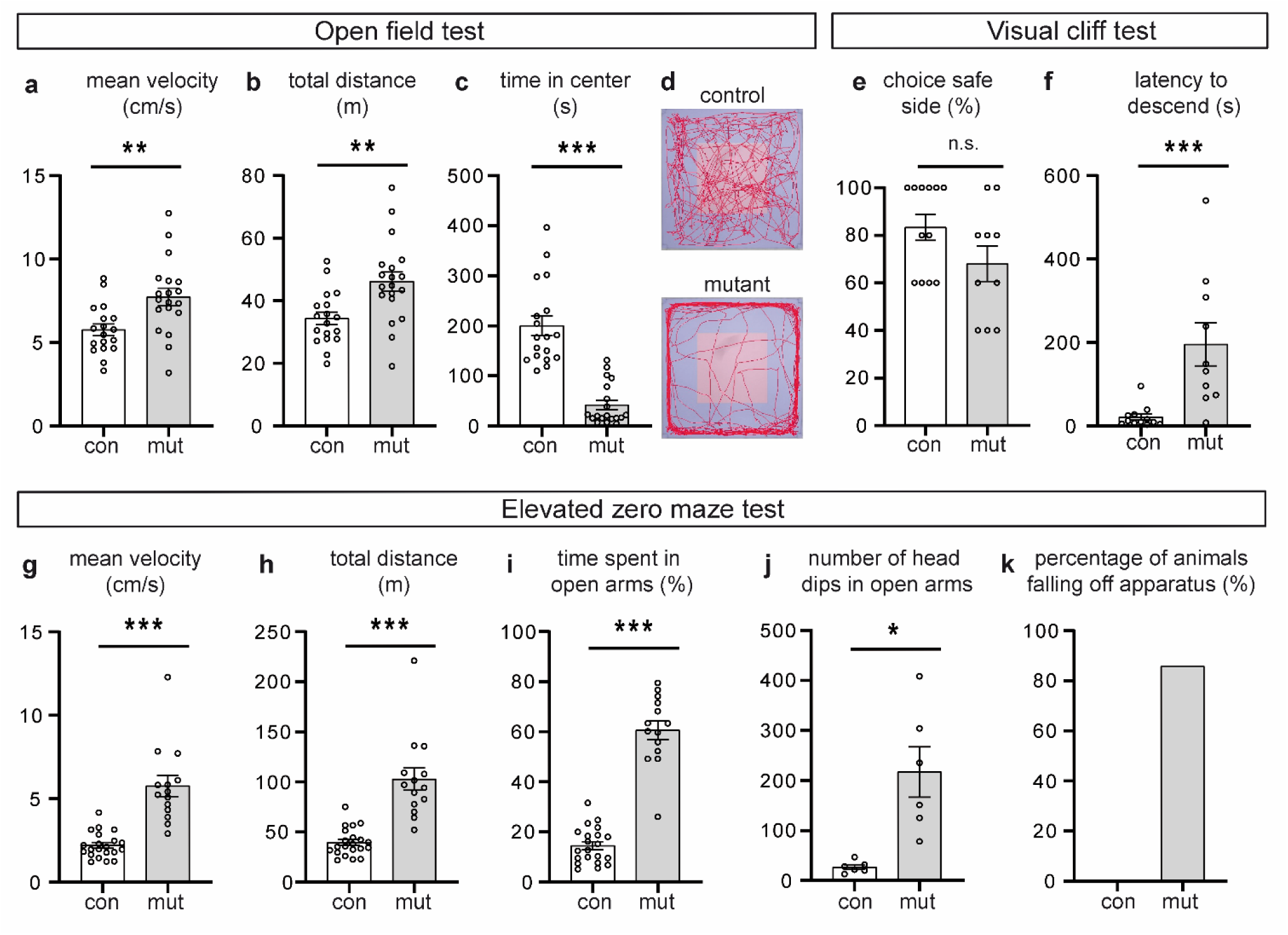
Anxiety assessment of adult control and mutant mice by open field and elevated zero maze tests, and visual cliff tests**. a**-**d** Open field test: mean velocity (**a**), total covered distance (**b**), and time spent in the centre (**c**). **d** represents typical track visualizations of single open field experiments. n =18 (control), n=19 (mutant). **e**, **f** Visual cliff test: percentage of attempts of mice choosing the “safe” side (**j**), latency to descend from the platform (**k**). n =12 (control), n=10 (mutant). **g-k** Elevated zero maze test: mean velocity (**g**), total covered distance (**h**), percentage of time spent in the open arms (**i**), number of head dips in the open arms (**j**) and percentage of animals falling off the apparatus (**k**). For **g**-**i, k** n =21 (control), n=14 (mutant); for **j**, n =6 (control), n=6 (mutant). Error bars indicate standard error of the mean. * *p* ≤ 0.05, ** *p* ≤ 0.01, *** *p* ≤ 0.001, n.s. – non-significant.

Increased locomotor activity in a novel environment, coupled with decreased time in the centre of an open field and paradoxically increased open arm exploration has also been reported in a Neurogranin knockout mouse model [56] and in mouse models of Alzheimeŕs disease [27, 44, 49]. In case of the former case, these findings were interpreted as anxiety-like behaviour associated with increased reactivity to novelty, panic-like behaviour, or elevated impulsivity. In the latter mouse models, this was judged as behavioural disinhibition, i.e. a poor self-regulation or the inability to control one’s activity level, attention, and emotions. To assess whether the increased head dipping frequency of mutant mice in the elevated zero maze test and beam walking test could be caused by impaired depth perception or visual acuity, we performed a visual cliff test. In this test, mice were placed on a narrow bar that separates two zones of a plexiglass plate: a “flat, seemingly safe side” with a checkerboard pattern is applied directly under the glass and a “deep side” where the same pattern is placed 60 cm below the glass, creating the illusion of a slope or cliff. Zone preference and the time taken to enter one of the zones were measured. Although there was a trend towards higher variability in zone preference among the mutant group, these animals chose the safe zone slightly less frequently than control animals (Fig. 7e), and no overall significant difference was obtained between the mutant and control animals (chi-square test, p-value = 0.4939087). This suggests that the increased frequency of head dipping detected in the elevated zero maze and beam walking tests is not associated with an impaired depth perception or visual acuity. Noteworthy, mutant mice required significantly more time to descend from the bar, i.e., to choose between the two sides (Fig. 7f). While on the bar, they walked back and forth, making head dips towards both safe and deep sides, without exhibiting signs of fear or agitation, such as freezing, scurrying, or adopting a stretch-attend posture. This behaviour has been interpreted as impaired decision-making or a slowed response capacity [25, 42].

Considering the outcomes of the open field, elevated zero maze, beam walking and visual acuity tests collectively, the behaviour of the mutant animals may be characterized as anxiety-like and agitation in response to unfamiliar environments. Furthermore, these findings could indicate deficits in decision-making, which could impair risk assessment abilities. This is evidenced by extreme vertical head dips observed in the elevated zero maze and beam walking tests, frequently resulting in falls from the apparatus.

### Individuals with *SMARCB1* pathogenic variants exhibit cognitive and behavioural abnormalities

Based on the behavioural findings in *Smarcb1* mutant mice, we sought to determine whether similar features are present in humans with SMARCB1-related neurodevelopmental disorders. We assessed general cognitive abilities using standardized intelligence testing in four individuals, and in three of them, additional cognitive and behavioural functioning was investigated (Supplementary Table 2). Parents of the three children completed the Child Behaviour Checklist (CBCL) to assess a broad range of emotional and behavioural problems, and the Behaviour Rating Inventory of Executive Function (BRIEF-2) to assess everyday executive functioning, including cognitive flexibility, impulse control, and working memory. Furthermore, the Strengths and Difficulties Questionnaire (SDQ) was used to evaluate emotional symptoms, conduct problems, hyperactivity/inattention, and prosocial behaviour. The developmental quotients (DQ) derived from the intelligence testing of the four individuals varied substantially, ranging from very low to average compared to same-aged peers (15-81). Within executive functioning, the self-monitor and plan/organize scales reached clinical scores in two of the two assessed individuals for these features. These domains reflect the ability to monitor and adjust one’s own behaviour, as well as to set goals, anticipate steps, and manage thoughts and materials to reach targets – skills that are highly relevant for effective decision-making as well as for managing behavioural operations. In two of the three assessed individuals, social behavioural problems and hyperactive/impulsivity problems were present. Thus, these results correspond to the findings obtained with the *Smarcb1* mutant mice regarding cognitive impairments and distinct behavioural abnormalities.

## Discussion

Heterozygous *SMARCB1* pathogenic variants have been identified in Coffin-Siris syndrome and ID-CPH. However, the relationship between *SMARCB1* and brain function is poorly understood, and there is only very limited information regarding the behavioural abnormalities associated with this gene. The presented results, obtained with a mutant *Smarcb1* mouse model and from behavioural assessments of individuals with *SMARCB1*-related neurodevelopmental disorders, demonstrate for the first time the implications of this gene for dendrite development and certain behavioural, cognitive, and motor skills. Furthermore, our findings imply a connection between a chromatin remodelling complex in the nucleus, neuronal structure changes, and a key regulator of actin polymerization. This offers a fresh perspective on BAF complexes and their associated brain pathologies.

### Functional disabilities in *Smarcb1* mutant mice and *SMARCB1*-related Coffin-Siris syndrome

Intellectual disability, delayed to absent expressive language development, delayed motor development, and hypotonia are commonly reported functional abnormalities in Coffin-Siris syndrome, and some studies describe different types and degrees of additional behavioural difficulties including hyperactivity, attention problems, social problems, and anxiety. However, there is a general lack of quantitative, comparable data, and among all investigated Coffin-Siris syndrome cases, very few functional assessments of individuals with *SMARCB1* pathogenic variants have been published (references in [43]; [77]). The studies that included probands with verified *SMARCB1* pathogenic variants generally reported developmental delay/intellectual disability, speech impairment, delayed motor development - the most severe being absence of walking ability - hypotonia, and behavioural abnormalities with hyperactivity being the most frequent one (three of eight cases) [41, 77]. These pathogenic variants are all located in exons 8 and 9, which encode the C-terminal domain of *SMARCB1*. There is one exceptional case of an individual with a *de novo* heterozygous c.568C > T (p.Arg190Trp) variant in the *SMARCB1* gene and normal IQ and adaptive functioning [6]. Individuals with another neurodevelopmental disorder associated with this gene, *SMARCB1*-related severe intellectual disability with choroid plexus hyperplasia (ID-CPH), have in general terms been described as exhibiting speech and motor delay in addition to intellectual impairment [18].

Regarding mouse models that mimic the genetic defects found in Coffin-Siris syndrome, behavioural analyses have also been performed on *Arid1b^+/-^* mice [14, 21, 35, 72]. Although some of the results obtained by four different groups using independently generated mouse lines deviate from each other, the phenotypes that all of the groups reported for *Arid1^+/-^*, and which are also present in the *Smarcb1* mutant mice are motor deficits, albeit based on different tests: reduced grip strength indicating muscle weakness [14, 21, 72], reduced locomotor activity in an open field [21, 35, 72] (and our study), reduced speed and increased food slips in beam walking tests [72] (and our study), and lower latency to fall in rotarod tests [35, 72]. Gait analyses of *Arid1^+/-^* mice in one study revealed normal motor coordination [21], which is consistent with our results on *Smarcb1* mutant mice. Another finding shared by the *Smarcb1* and three of these *Arid1b* mouse models are anxiety-like phenotypes. The cognitive impairment present in *Smarcb1* mutant mice, hippocampal-dependent spatial learning and memory, has also been reported by two studies on *Arid1b^+/-^* mice [35, 72], however results obtained by the other two groups do not support any cognitive deficits [14, 21].

The abnormalities we found in *Smarcb1* mutant mice are in accordance with the main functional disability categories generally described for Coffin-Siris syndrome and for *SMARCB1*-related ID-CPH. Impaired spatial learning and memory of the mutant mice fits with the intellectual disability found in these two human conditions.

Quantitative data on cognitive function such as DQ or intelligence quotient (IQ) have not been reported for *SMARCB1*-related Coffin-Siris syndrome associated with the classical C-terminal alterations. Among the four individuals examined with *SMARCB1* pathogenic variants in this study, DQs varied widely, even between the two children C and D with identical variants (Supplementary Table 2). Developmental delay in gross motor skills (e.g., sitting, walking), and hypotonia as well as fine motor skills (e.g., difficulties with tasks requiring finger coordination) have been reported for Coffin-Siris syndrome [10, 70, 76]. The third behavioural conspicuity, anxiety-like behaviour and agitation in an unfamiliar environment, have not yet been recognized as typical for Coffin-Siris syndrome or *SMARCB1*-related ID-CPH. However, hyperactivity is a behavioural characteristic that resembles one of the features commonly seen in attention deficit hyperactivity disorder (ADHD), as reported in some Coffin-Siris syndrome cases [41, 67–70] including 3 of 15 with *SMARBC1* pathogenic variants [41, 67]. Among the four assessed children with *SMARCB1* pathogenic variants and Coffin-Siris syndrome, the one with the highest developmental quotient (81.54) exhibited anxious behaviour (Supplementary Table 2). Given the small sample size and wide range in general cognitive functioning, it remains unclear whether children with lower DQ scores display fewer anxiety-related behaviours because they are genuinely less pronounced, expressed differently, or less easily observed. Among the three children whose investigations included hyperactivity/impulsivity scores (individuals A, B, and D), one showed clear behavioural signs of hyperactivity/impulsivity, one showed mild difficulties, and one did not display noticeable problems (Supplementary Table 2). Notably, our data suggest an additional, hitherto unreported behavioural aspect, which is based on tests of the *Smarcb1* mutant mice and hinted at in the investigations of patients, namely, impaired decision-making. Two individuals (A and B) who were assessed for executive functions in everyday functioning (BRIEF-2), showed clear difficulties in self-monitoring and planning and organizing, which may contribute to the observed impairments in decision-making. As impaired decision-making leads to an inaccurate risk assessment, this might be highly relevant in a clinical setting and should be considered in human studies and care.

### Altered chromatin remodelling complex components lead to dendritic defects

Dendritic defects, such as reduced dendrite branching, length, and number, are known features of several intellectual disability disorders [62]; however, to our knowledge dendrite abnormalities have so far not been investigated in *postmortem* Coffin-Siris syndrome brain tissue. The dendritic defects in *Smarcb1* mutant mice reported in this study resemble findings in murine cells with reduced or absent expression of two other genes associated with Coffin-Siris syndrome: *Arid1b* and *Smarca4*. *Arid1b* knockdown in mouse brain tissue upon *in utero* gene delivery leads to reduced dendritic arborization in primary cortical and hippocampal neuron cultures [36]. Deletion of *Smarca4* results in reduced dendrite length of P7 cortical pyramidal cells and absent P7 Purkinje cell dendrites in *GFAP-Cre Brg1^fl/fl^* mice [34]. Reduced axon and dendrite numbers of cortical pyramidal neurons are found in Nex-Cre *Brg1^fl/fl^* animals [17]. Moreover, knockout or knockdown of other genes encoding components of ATP-dependent chromatin remodelling complexes is linked to dendritic defects in mouse brains and cultured cells. These code for the BAF complex components BAF53B [9, 81] and BCL7a [80], for the catalytic subunit of ISWI family complexes SMARCA5 [4], and for CHD4 and CHD8 proteins of the CHD family of chromatin remodelling complexes [19, 83, 84] (summarized in Supplementary Table 1). Finally, expression reduction of the *SMARCB1* homolog in Drosophila, *Snr1*, is associated with increased primary dendrite extension and decreased lateral branching of sensory neurons [58].

Both intrinsic and extrinsic factors regulate dendritic development. During the first postnatal week of mouse development, intrinsic factors are assumed to play a key role in Purkinje cells [20]. The complex and highly regulated process of dendrite development requires precise and temporally fine-tuned control of gene expression and chromatin remodelling complexes. Our findings and dendritic abnormalities reported for mouse tissue of two other Coffin-Siris genes, *Smarca4* and *Arid1b,* point to a cell-intrinsic function of these BAF complex genes on dendrite development: We detected dendritic defects at a very early stage, in brain tissue from P7 mutant mice, a stage at which also *Smarca4* mutant animals show dendrite abnormalities. In addition, knockdown of *Arid1b* negatively impacts dendritic development in cultured neurons outside their normal tissue context [36]. In addition, this interplay with extrinsic factors, including growth factors and neuronal activity influenced by synaptic contacts, likely impacts BAF complex-dependent dendrite development *in vivo*. Cerebellar hypoplasia in mutant mice, which is already present before birth, suggests that a delayed brain maturation could contribute to the impaired dendritic development. Moreover, cultured primary hippocampal neurons from mice lacking *Baf53b* show a severe reduction in activity-dependent dendritic outgrowth [81] and neuronal activity influences nBAF assembly, BAF composition, and ATPase activity, thereby mediating rapid transcription and chromatin accessibility [16, 29]). Notably, our finding of reduced *Wasl* transcripts in mutant Purkinje cell nuclei suggests that an impaired regulation of actin polymerization may be involved in dendritic defects in a BAFopathy. *Wasl*, which encodes N-WASP, is broadly expressed, including neurons, and promotes the nucleation of a new actin filament from an existing one by the ARP2/3 complex, leading to actin filament branches [13, 50, 53]. In neurons, branched actin structures are present and regulated by N-WASP and ARP2/3 at multiple locations: in growth cones at the tips of extending neurites or axons, where they drive membrane protrusion and directional navigation, at dendritic branch initiation sites, and in dendritic spines where they are crucial for spine formation and morphology [40, 57, 78]. In addition, N-WASP also regulates ARP2/3-mediated actin polymerization in the nucleus, thereby impacting gene transcription [79, 82, 85]. Interestingly, WASP, the founding member of the protein family, has been shown to associate with BAF complex components including SMARCB1 (designated BAF47 in the respective publication) [66]. Our finding of reduced *Wasl* expression in *Smarcb1* mutant Purkinje cell nuclei, coupled with the integral role of nuclear actin and actin-related proteins in BAF complexes, suggests further investigation into the relationships between WASP in the context of BAFopathies, including its dual cytoplasmic and nuclear localizations as well as its function.

### Dendritic tree alterations and brain function

Establishing a causal relationship between alterations in dendrite length or dendritic tree complexity and neuronal network or brain function *in vivo* remains a significant challenge within neuroscience. This difficulty arises primarily from the technical complexities involved in simultaneously manipulating dendrites and obtaining corresponding readouts at these levels. Our assessment of both morphological and behavioural alterations within the same mouse model, alongside comparison of mouse and human behavioural findings, provides evidence linking morphological changes to behavioural outcomes, enhancing our understanding of the functional consequences of SMARCB1 gene alterations. Furthermore, the field benefits from experimental results obtained from simpler organisms, computer simulations, and artificial neural networks. It is important to note that, in addition to transmitting input signals to the neuronal cell body, dendrites possess computational capabilities themselves [60]. Notably, introducing key dendrite features – their branched structure, active ionic mechanisms and synaptic plasticity – into artificial neural networks lowers their computational cost of solving a problem, thus increasing their efficiency [15]. Including dendrites in a model of a neuronal network in the mouse cortex in the context of a decision-making task increased neuronal selectivity, leading to a reduction in error trials [5]. The selective abolishment of dendrites in *Drosophila* motor neurons demonstrated that a reduced dendritic architecture is dispensable for basic motor performance, but leads to performance deficits in sophisticated motor tasks and mating-related motor behaviour [64]. Simulating dendritic reduction in digitally reconstructed neurons revealed a decrease in dynamic range, i.e. the range of input that a neuron can encode in its output, and thus distinguish. These results were proposed as a neural basis for the reduced ability of older persons to distinguish between similar experiences (pattern separation) as the complexity and size of the dendritic arbor is reduced with ageing [39]. Concerning the compartmentalization of dendritic activity, decreased branch numbers could also lead to an impaired performance in decision-making, since different levels of calcium signals in single branches of dendrites are associated with a decision making task [38]. Taken together, the reduced number of dendritic branches and the reduced length of dendrites in neurons, as observed in the *Smarcb1* mutant mouse model, could result in fewer receptive connections from other neurons. This could lead to decreased information integration and input discrimination in neurons, as well as lower computational efficiency in neuronal networks. Based on the aforementioned published data, these alterations could underlie deficits in complex brain functions, such as learning, memory, decision-making in *Smarcb1* mutant mice and individuals with *SMARCB1*-related neurodevelopmental disorders.

Finally, we would like to highlight the potential therapeutic implications of our findings. Even long after the establishment of dendritic trees in adult mice, reduced dendritic length and complexity as well as impaired sensory-motor functions can be significantly improved by correcting the underlying genetic cause as demonstrated in a mouse model of the severe developmental disorder Rett syndrome [63]. Given the monogenic cause of *SMARCB1*-associated neurodevelopmental disorders and based on our results and the dendritic changes associated with defects in other components of chromatin remodelling complexes, targeting the corresponding genes or proteins could therefore be considered a potential therapeutic entry point.

## Supplementary Information

The online version contains supplementary information available at *Acta Neuropathologica* online.

## Data availability

All data used and analysed in this study are available from the corresponding author upon reasonable request.

## Declarations

### Ethics approval and consent to participate

The patients’ consent was obtained in accordance with the Declaration of Helsinki and approved by the Medical Ethical Commission of the East of the Netherlands (2021-7478). All animal procedures were performed in compliance with German national guidelines and approved by the local authorities (approval for the conduct of scientific experiments DA8 / 2000, Regierungspräsidium Darmstadt, Hesse, Germany).

### Consent for publication

Consent for publication was not required as all data were fully anonymized and no identifiable information is included.

## Acknowledgements

We are grateful to Meike Stotz-Reimers and Liliana Davkova for excellent technical support.

## Author contributions

All authors contributed to the study conception and design. Material preparation, data collection and analysis were performed by Kristina I. Lemke, Alina Filatova, Joanna Chiang, Hannah North, Myrthe R. M. Kamphof, and Michaela Becker-Röck. Figures and tables were generated by Kristina I. Lemke, Alina Filatova, Joanna Chiang, Hannah North, Myrthe R. M. Kamphof, Michaela Becker-Röck, Hanna Swaab, and Gijs W. E. Santen. The first draft of the manuscript was written by Ulrike A. Nuber, and all authors contributed to its development, reviewed the text, and approved the final version.

