## Supplementary material for "Chromatin remodelling subunit SMARCB1 is implicated in dendrite development and complex brain functions"

Correspondence to: Professor Ulrike A. Nuber

### Supplementary Tables

**Supplementary Table 1** Chromatin remodelling complex components associated with dendritic tree abnormalities. Arrows indicate reduced or increased dendrite length and dendritic tree complexity/number.

ATPase subunits are depicted in orange, BAF base module subunits in blue, and BAF ARP module subunits in green. Gray subunit of ISWI complex illustrate an interchangeable subunit. In case of CHD, grey subunits are part of the NuRD complex, which contains CHD4/5.

**M = mouse, H = human, u = untreated, d = upon depolarization, * = this study**

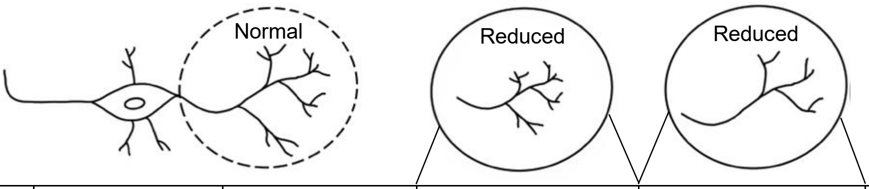


| Complex | Subunit | Module | Dendrite length | | Dendritic tree complexity/ Dendrite number | | Species | Ref. |
|---|---|---|---|---|---|---|---|---|
| | | | *In vivo* | *In vitro* | *In vivo* | *In vitro* | | |
| 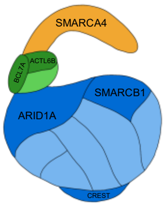  BAF | **ARID1B** | **Base** | ↓ | ↓ | ↓ | ↓ | **M** | **[16]** |
| | **ACTL6B** | **ARP** | | ↓ **u, d** | ↓ | ↓ **u, d** | **M** | **[31]** |
| | | | | | | ↓ | **H** | **[5]** |
| | **BCL7A** | **ARP** | | | ↓ | | **M** | **[30]** |
| | **CREST** | **Base** | ↓ | ↓ **d** | ↓ | | **M** | **[2]** |
| | **SMARCA4** | **ATPase** | | | ↓ | | **M** | **[7]** |
| | | | ↓ | | | | **M** | **[14]** |
| | **SMARCB1** | **Base** | ↓ | | ↓ | | **M** | * |
| 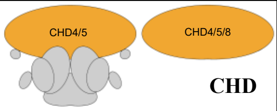  CHD | **CHD4** | **ATPase** | ↑ | | ↑ | | **M** | **[33]** |
| | **CHD5** | **ATPase** | | | | ↓ | **M** | **[22]** |
| | **CHD8** | **ATPase** | | | ↓ | | **M** | **[8]** |
| | | | ↓ | ↓ | ↓ | ↓ | **M** | **[32]** |
| 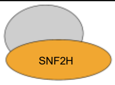  ISWI | **SNF2H** | **ATPase** | ↓ | | | | **M** | **[3]** |

**Supplementary Table 2** Results of questionnaires and general cognitive functioning (DQ) in children with *SMARCB1* pathogenic variants.

| | **Individual A** | **Individual B** | **Individual C** | **Individual D** |
|---|---|---|---|---|
| **Pathogenic variant** | c.1120C>T p.Arg374Trp | c.1089G>T, p.Lys363Asn | c.1091_1093del p.Lys364del | c.1091_1093del p.Lys364del |
| **Age, Sex** | 13 years, boy | 10 years, boy | 13 years, girl | 5 years, girl |
| **Independent walking age (months)** | 14 | 26 | - | 16 |
| **Age first words (months)** | 24 | 60 | - | Speaks three words |
| **Tube feeding** | Yes, since 2 months | No | Since birth | Yes, 4 weeks after birth and again from 18 months- 4 years. |
| **Epilepsy** | No | No | No | No |
| **MRI** | Normal | Underdeveloped corpus callosum, large fossa posterior with mega cisterna magna, large 4th ventricle, minor rotation of vermis | Megacisterna magna + cerebellar hypoplasia | N/A |
| **Developmental Quotient (DQ)** | 81.54 | 43.50 | 15.09 | 40.32 |
| **Child Behaviour Checklist (CBCL 1,5-5 or 6-18)** | | | | |
| **Total problem behaviour** | ++ | - | N/A | - |
| **Internalizing behaviour** | + | - | N/A | - |
| **Externalizing behaviour** | ++ | - | N/A | - |
| **Syndrome scale scores 1,5-5 or 6-18** | | | | |

| | | | | |
|---|---|---|---|---|
| **Anxious/depressed** | ++ | - | N/A | - |
| **Withdrawn/ depressed** | + | - | N/A | - |
| **Somatic complaints** | ++ | - | N/A | - |
| **Social Problems** | ++ | - | N/A | N/A |
| **Thought Problems** | ++ | - | N/A | N/A |
| **Attention Problems** | + | - | N/A | - |
| **Rule-Breaking Behaviour** | - | - | N/A | N/A |
| **Aggressive Behaviour** | ++ | - | N/A | - |
| **Behaviour Rating Inventory of Executive Function (BRIEF-2 (T-scores))** | | | | |
| **Inhibit** | ++ (T=65) | + (T=60) | N/A | N/A |
| **Self-monitor** | ++ (T=66) | ++ (T=69) | N/A | N/A |
| **Shift** | + (T=60) | + (T=62) | N/A | N/A |
| **Emotional Control** | + (T=62) | - (T=52) | N/A | N/A |
| **Initiate** | - (T =54) | ++ (T=72) | N/A | N/A |
| **Working Memory** | - (T=58) | ++ (T=78) | N/A | N/A |
| **Plan/Organize** | ++ (T=65) | ++ (T=70) | N/A | N/A |
| **Task-Monitor** | - (T=54) | ++ (T=70) | N/A | N/A |
| **Behavioural Regulation Index** | - (T=54) | ++ (T=71) | N/A | N/A |
| **Emotional Regulation Index** | ++ (T=74) | - (T=56) | N/A | N/A |
| **Cognitive Regulation Index** | + (T=62) | ++ (T=75) | N/A | N/A |

| | | | | |
|---|---|---|---|---|
| **Global Executive Composite** | - (T=57) | ++ (T=71) | N/A | N/A |
| **Strengths and Difficulties Questionnaire (SDQ)** | | | | |
| **Emotional problems** | ++ | - | N/A | - |
| **Behavioural problems** | ++ | + | N/A | - |
| **Hyperactivity/ impulsivity** | ++ | + | N/A | - |
| **Peer problems** | + | ++ | N/A | - |
| **Pro-social behaviour** | +++ | +++ | N/A | - |
| **Total difficulties** | +++ | ++ | N/A | - |

**Table legend:** N/A = not available; ++ = clinical scores, values in the higher range, suggesting significant difficulties or impairments; + = borderline scores, indicates moderate difficulties and may require monitoring or further evaluation; - = non-clinical scores, indicating typical or no significant behavioural or cognitive difficulties. *SMARCB1* transcript used NM_003073.3.

### Supplementary Figures

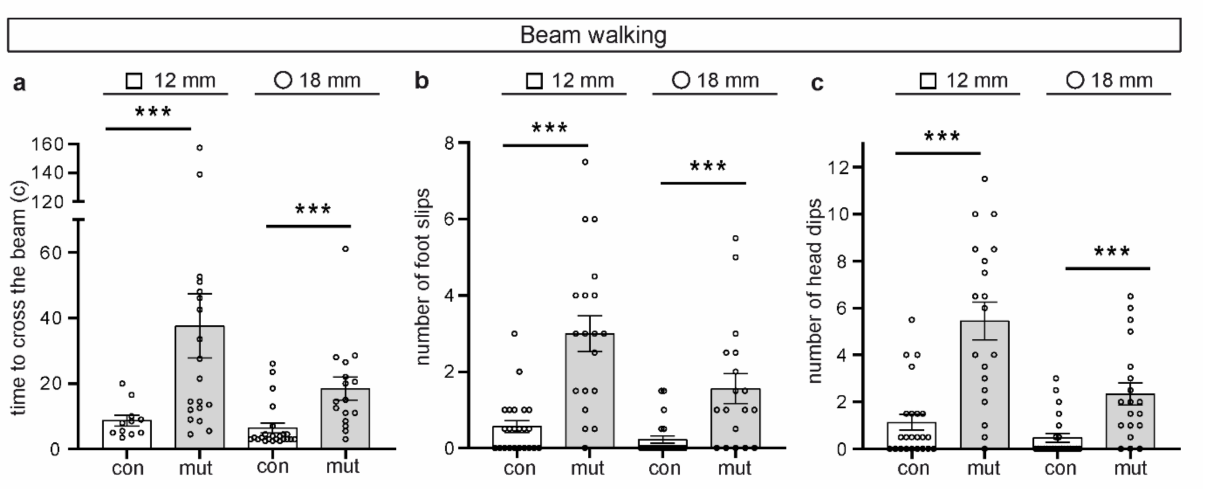


**Supplementary Figure 1** Beam walking test using 12 mm square and 18 mm round beams. **a** Time to cross the beam, n=11 (control, 12 mm square beam), n=23 (control, 18 mm round beam), n=19 (mutant, 12 mm square beam), n=15 (mutant, 18 mm round beam). **b** Number of foot slips, n=23 (control), n=19 (mutant). **c** Head dips, n=23 (control), n=19 (mutant). * $p \leq 0.05$, ** $p \leq 0.01$, *** $p \leq 0.001$, n.s. – non-significant.

#### Morris water maze test

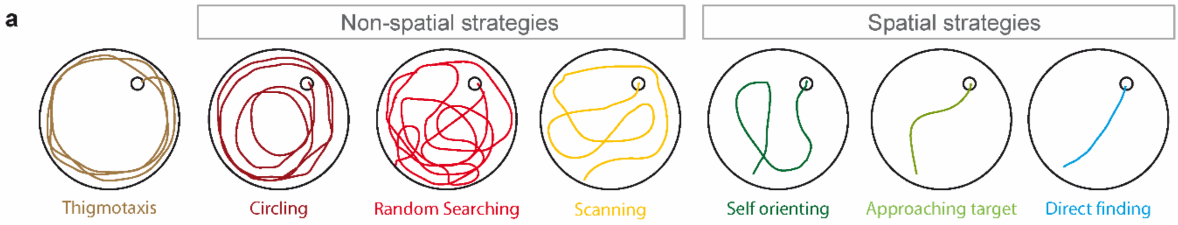


**b** Initial acquisition

| | con 1 | con 2 | con 3 | con 4 | con 5 | con 6 | mut 1 | mut 2 | mut 3 | mut 4 | mut 5 | mut 6 | mut 7 |
|---|---|---|---|---|---|---|---|---|---|---|---|---|---|
| Day 1 | 17 | 7 | 15 | n.r. | 7 | 23 | n.r. | 8 | n.r. | n.r. | 80 | 18 | n.r. |
| | n.r. | 43 | 18 | n.r. | 77 | 35 | 17 | 22 | 10 | 110 | n.r. | 45 | 107 |
| | 19 | 12 | 14 | 10 | 104 | 17 | 60 | n.r. | n.r. | n.r. | 3 | 3 | n.r. |
| | 11 | 53 | 7 | 2 | 22 | 3 | 18 | 9 | 32 | n.r. | 24 | 41 | 12 |
| Day 2 | 29 | 32 | 5 | 23 | 37 | 8 | n.r. | n.r. | 44 | 20 | 16 | 24 | 8 |
| | 33 | 6 | 4 | 17 | 26 | 10 | 44 | 25 | n.r. | 27 | 55 | 4 | 56 |
| | 31 | 3 | 19 | 23 | 17 | 5 | 4 | 24 | 33 | n.r. | 12 | 3 | 12 |
| | 21 | 9 | 11 | 34 | 4 | 27 | 6 | 9 | n.r. | n.r. | 12 | 12 | 64 |
| Day 3 | 8 | 12 | 21 | 6 | 16 | 19 | n.r. | n.r. | 88 | 7 | 76 | 5 | 24 |
| | 3 | 2 | 9 | 2 | 3 | 2 | n.r. | 2 | n.r. | n.r. | 47 | 5 | 31 |
| | 6 | 8 | 6 | 10 | 7 | 7 | 5 | n.r. | 20 | n.r. | 4 | 5 | 53 |
| | 7 | 7 | 12 | 3 | 5 | 15 | 14 | 46 | 20 | 18 | 6 | 9 | 29 |
| Day 4 | 5 | 6 | 7 | 13 | 9 | 16 | 11 | 24 | 13 | 89 | n.r. | 9 | 7 |
| | 17 | 3 | 7 | 19 | 4 | 25 | n.r. | 15 | 55 | n.r. | 4 | 6 | 37 |
| | 8 | 4 | 10 | 11 | 4 | 8 | 23 | 12 | 14 | n.r. | 13 | 9 | 5 |
| | 3 | 9 | 13 | 3 | 3 | 7 | 107 | 39 | 13 | 106 | 3 | 21 | 29 |
| Day 5 | 8 | 3 | 31 | 39 | 4 | 9 | 28 | 46 | 17 | 59 | 8 | 12 | 26 |
| | 7 | 19 | 15 | 8 | 4 | 8 | 108 | 5 | 57 | 6 | 15 | 30 | 2 |
| | 4 | 8 | 30 | 25 | 3 | 27 | 12 | 26 | 16 | 5 | 111 | 3 | 14 |
| | 7 | 9 | 29 | 9 | 2 | 20 | 12 | 7 | 7 | 119 | 94 | 58 | 3 |

**c** Reversal learning

| | con 1 | con 2 | con 3 | con 4 | con 5 | con 6 | mut 1 | mut 2 | mut 3 | mut 4 | mut 5 | mut 6 | mut 7 |
|---|---|---|---|---|---|---|---|---|---|---|---|---|---|
| Day 1 | 60 | 26 | 7 | 28 | 41 | 22 | 66 | 120 | 103 | 116 | n.r. | 60 | n.r. |
| | 7 | 16 | 32 | 14 | 21 | 7 | 90 | 5 | 48 | n.r. | 5 | 101 | 13 |
| | 7 | 40 | 10 | 17 | 13 | 33 | 60 | 51 | 8 | n.r. | 5 | 26 | 12 |
| | 12 | 5 | 22 | 7 | 12 | 35 | 22 | 10 | 13 | 9 | 18 | 3 | 6 |
| Day 2 | 6 | 5 | 21 | 44 | 14 | 12 | 31 | 46 | 5 | 108 | 61 | 30 | 13 |
| | 19 | 3 | 20 | 30 | 22 | 16 | 77 | 84 | n.r. | 26 | 31 | 24 | 37 |
| | 7 | 9 | 15 | 8 | 11 | 28 | 9 | 49 | 10 | 86 | 101 | 9 | 31 |
| | 10 | 4 | 13 | 24 | 8 | 12 | 64 | 50 | 47 | 98 | 61 | 5 | 14 |
| Day 3 | 9 | 4 | 4 | 19 | 8 | 12 | 30 | 52 | n.r. | 23 | 26 | 13 | 10 |
| | 3 | 6 | 8 | 4 | 6 | 5 | 27 | 43 | 116 | 13 | 11 | 12 | 15 |
| | 4 | 15 | 9 | 10 | 7 | 9 | 9 | 18 | 119 | n.r. | 94 | 12 | 11 |
| | 9 | 8 | n.r. | n.r. | 5 | 8 | 28 | 7 | 13 | 69 | 14 | 10 | 17 |
| Day 4 | 10 | 9 | 15 | 5 | 8 | 5 | 76 | 12 | n.r. | 14 | 10 | 8 | 19 |
| | 8 | 10 | 13 | 12 | 2 | 11 | 16 | 73 | n.r. | 10 | 3 | 9 | 11 |
| | 5 | 3 | 11 | 6 | 5 | 3 | 14 | 7 | n.r. | 5 | 8 | 17 | 13 |
| | 18 | 2 | 4 | 11 | 8 | 4 | 42 | 58 | n.r. | 4 | 7 | 8 | 12 |
| Day 5 | 20 | 7 | 13 | 15 | 4 | 13 | 11 | 43 | n.r. | 25 | 47 | 6 | 10 |
| | 3 | 3 | 4 | 7 | 5 | 8 | 86 | 88 | n.r. | 25 | 6 | 10 | 15 |
| | 5 | 2 | 3 | 10 | 3 | 5 | 9 | 56 | n.r. | 24 | 54 | 10 | 20 |
| | 4 | 3 | 21 | 13 | 13 | 8 | 11 | 54 | n.r. | 31 | 61 | 3 | 3 |

**Supplementary Figure 2** Exploration strategies employed by adult control and mutant mice. **a** The applied exploration strategy classification with characteristic swim paths: thigmotaxis, non-spatial strategies, and cognitively sophisticated spatial strategies (employing spatial cues). **b**, **c** The dominating exploration strategies of single control and mutant animals on days 1-5 of initial acquisition (**b**) and reversal learning (**c**) (four trials per day). The times (in seconds) animals took to reach the platform at each single trial are indicated; n.r. indicates that the animal did not reach the platform within the maximum trial time of 120s.

### Supplementary Methods

#### Patient consent

The subjects' consent was obtained according to the Declaration of Helsinki and been approved by the Medical Ethical Commission of the East of the Netherlands (2021-7478).

#### Animals

*Smarcb1*$^{+/inv}$ *NesCre*$^{+/-}$ (mutant) and *Smarcb1*$^{+/inv}$ *NesCre*$^{-/-}$ (control) mice were described previously (Filatova et al. 2019). All animal procedures were performed in accordance with national guidelines and approved by the local authorities (approval for the conduct of scientific experiments DA8 / 2000, Regierungspräsidium Darmstadt, Hesse, Germany). The corresponding ethical application was prepared before the study and included the research question, key design features, analysis plan, as well as the maximal number of animals for the various experiments, the endpoints established for the study, the signs that were monitored, and the frequency of monitoring. The animals were housed under a constant temperature (22 ± 1°C), humidity and an automatically controlled 12-h light-dark cycle (lights on at 6 a.m.), with access to food and water ad libitum. For behavioural tests, animals with milder phenotypes, i.e., that had reached age of 10 weeks, were used. All behavioural tests were performed with female and male mice aged 12-26 weeks. Whole embryos were isolated at E16.5; mouse brains were isolated between P7 and P32. No animals have been excluded from the analyses.

#### Staining of tissue sections

Whole embryos (E16.5) or mouse brains were isolated and dissected brain tissue was fixed in 4% paraformaldehyde (PFA) in phosphate-buffered saline (PBS) for different durations of time depending on the developmental stage. After two wash steps with PBS, dehydration, and paraffin embedding, brain tissue was cut into 5 µm thick sections using a Jung RM2055 device (Leica Biosystems) and transferred to Thermo Scientific Polysine Adhesion microscope slides. For dendritic analysis of Purkinje cells at P10, that were stained with calbindin antibody, 10µm sections were used. Prior to stainings, sections were deparaffinized and rehydrated.

For hematoxylin and eosin staining, slides were incubated in hematoxylin for 3–5 min, in tap water for 2 min, stained with eosin Y for 4 min, rinsed with water, dehydrated gradually in ethanol (70%, 80%, 90%, and 100%, 30 s each step), and mounted in Entellan. Images were acquired using an Aperio CS2 slide scanner (Leica Biosystems, RRID:SCR_025111).
For immunofluorescence staining, sections were incubated in epitope retrieval citrate buffer (10mM Sodium Citrate, 0.05% Tween 20, pH 6.0) for 40 minutes in a water bath heated to 100°C and subsequently cooled down for 40-60 minutes at room temperature. After a washing step in PBS, sections were permeabilized in 0.5% Triton X-100 in PBS for 10 minutes. Primary antibodies were applied in 0.01% Triton X-100/PBS overnight and incubated at 4°C and 0.01% Triton X-100/PBS only served as negative control. Sections were washed in PBS, followed by a one-hour incubation step with respective secondary antibodies diluted 1:300 in 0.01% Triton X-100 in PBS. Specimens were then incubated in 1 µg/mL 4′,6-diamidino-2-phenylindole (DAPI) solution (10236276001, Merck, Darmstadt, Germany) for 10 min, washed in PBS (5 min), rinsed briefly in water, dehydrated in ethanol, and air dried. For mounting, Fluorescence Mounting Medium (DAKO/Agilent) and coverslips (Thermo Scientific, Germany) were used. Immunofluorescent images were taken with a Zeiss Observer D1 microscope.

The following primary and secondary antibodies were used: anti-calbindin (D-28K) (chicken IgY polyclonal, Synaptic Systems, 214 006, 1:400, RRID:AB_2619903) and anti-BAF47/SMARCB1 (B-1) (rabbit monoclonal, Cell Signaling Technology, 91735, 1:800, RRID:AB_2800172); donkey anti-chicken AF488 (Dianova, #703-545-155, 1:300, RRID:AB_2340375) and donkey anti-rabbit Cy5 (Dianova, #711-175-152, 1:300, RRID:AB_2340607).

#### Golgi-Cox staining

Golgi-Cox staining was performed using the FD Rapid GolgiStain Kit (PK401A, FD NeuroTechnologies, Inc.) according to the manufacturer's instructions. Sagittal cerebellum sections and coronal forebrain sections from 32-week-old mice were investigated. Brain samples were impregnated for two weeks in the dark in a solution consisting of equal parts of solution A and solution B, and the solution was replaced 6h after tissue immersion. The brains were then transferred into solution C for cryoprotection and incubated for 72h; solution C was replaced once after 24h. After cryoprotection, the tissue samples were frozen in isopentane precooled with dry ice and stored at -80°C. The tissue was embedded in Tissue-Tek O.C.T. Compound, and 200 µm sections were generated at -16°C using a Thermo Scientific CryoStar NX50 cryostat and transferred onto gelatine-coated slides. After drying overnight, the sections were washed twice with distilled water at 4°C for 4 min each, incubated with staining solution consisting of equal parts of solution D and solution E for 10 min with gentle shaking and washed twice with distilled water at 4°C for 4 min each. Thereafter, the sections were dehydrated using ascending degrees of ethanol, defatted by washing twice with 100% xylene for 4 min each, and mounted using Entellan rapid mounting medium (#1.07960, Millipore) and stored in the dark before dendritic analysis.

#### Analysis of H&E stainings

Images of H&E-stained mouse cerebellum sections were analysed using the Aperio ImageScope software (pen tool function, RRID:SCR_020993) to measure the overall cerebellum section area and layer areas. Sections of three independent biological replicates (three animals per genotype) from five-months-old control and mutant mice were used.

#### Analysis of immunostaining fluorescence intensities

For fluorescence intensity measurements of P7, P10, and P21 immunostained tissue sections, ImageJ Fiji (RRID:SCR_002285) was used. 5-15 Purkinje and granule cells were chosen per section (*n*=3 animals per genotype, three technical replicates each) and the mean fluorescence of nuclei was measured.

#### Dendritic analyses of Purkinje cells at P7 and P10

5 µm and 10 µm paraffin sagittal sections of P7 and P10 brains, respectively, were stained with an anti-calbindin antibody as described above. Fifty Purkinje cells located on rostral and caudal sides of lobule VIII were analysed in control (*n*=3) and mutant (*n*=3) cerebelli using the ImageJ software (NIH, Bethesda, MD, USA [25], RRID:SCR_003070) and the fixed length line tool macro (Schmid and Rasband, 2015). Maximal dendritic length and width of primary dendrites at 4 µm (P7) or 8 µm (P10) apart from their soma origin were measured. For P7, the degree of dendritic branching was determined as the number of dendrites intersecting with a circle with a

diameter of 60 µm placed on the centre of the soma. For P10, the number of terminal dendritic ends was counted 100 µm along the Purkinje cell monolayer on three different sections per animal.

#### Dendritic analyses of adult neurons

200 µm paraffin sagittal cerebellum sections and coronal forebrain sections of control and mutant mice stained with the Golgi-Cox method were used to assess dendritic morphologies. To enable a detailed analysis of the neurons, Z-stack images (15 layers, 10 µm apart from each other) were taken of each section using the Aperio CS2 slide scanner (RRID:SCR_025111) and the program Aperio ScanScope (Leica Biosystems). The analysis was done using ImageJ software (NIH, Bethesda, MD [25], RRID:SCR_003070) and the fixed length line tool macro (Schmid and Rasband, 2015). Forty-nine Purkinje cells of adult control mice ($n$ = 5) and 42 Purkinje cells of adult mutant mice ($n$ = 8) were analysed. Maximal dendritic length, the width of primary dendrites at 10 µm apart from their soma origin, the number of terminal dendritic ends, and the degree of dendritic branching were analysed. Hippocampal and cortical neurons from 3 control and 7 mutant adult mice were analysed. For ten pyramidal neurons in the CA1 region, the maximal dendritic length, the width of primary apical dendrite at 10 µm apart from its origin in soma, and the number of primary basal branches were determined. For granule neurons in the upper blade of the dentate gyrus (n=10 for control, n=7 for mutant), the maximal dendritic length, the width of primary apical dendrites at 10 µm apart from their soma origin, and the degree of dendritic branching were assessed. For 20 pyramidal neurons in cortical layers III and V, the width of primary apical dendrites at 10 µm apart from their soma origin, the number of primary basal branches, and the degree of basal branching were measured. The degree of dendritic branching was determined as the number of dendrites intersecting with a circle with a diameter of 80 µm (pyramidal neurons in cortical layers III and V), 90 µm (Purkinje cells), or 100 µm (granule neurons located in the supra-pyramidal blade of the dentate gyrus) placed on the centre of the soma.

#### Nuclei isolation and fluorescence-activated nuclei sorting (FANS)

The protocol was adapted from Bartelt et al. [4]. Cerebelli were isolated from P7 mice and frozen in liquid nitrogen for at least 18h. 12-well plates and 50 ml conical tubes were coated with 1% BSA in dissociation buffer (82 mM $Na_2SO_4$, 10 mM glucose, 10 mM HEPES, 5 mM $MgCl_2$) for at least 1h at 4°C. Cerebelli were transferred into wells of 12-well plates containing 2 ml of extraction buffer (82 mM $Na_2SO_4$, 10 mM glucose, 10 mM HEPES, 5 mM $MgCl_2$, 1% Poly(1-vinylpyrrolidone-co-vinyl acetate), 1% Triton X-100, 0.01% BSA, 200 U RNase inhibitor). Tissue was triturated by pipetting up and down 20 times with a cut-off 1 ml pipette tip and 15 more times with a normal 1ml pipette tip. Following dissociation, samples were passed twice through a 25-gauge needle and filtered into a 50 ml conical tube through a 40 µm cell strainer. The filter was rinsed with 900 µl DAPI solution followed by a 1-minute incubation on ice. 20 ml wash buffer (82 mM Na2SO4,10 mM glucose, 10 mM HEPES, 5 mM $MgCl_2$, 0.01% BSA, 6.6 U RNase inhibitor) were added and samples were centrifuged at 850g at 4°C for at least 10 minutes until a stable pellet appeared. Pellets from three to four control and mutant samples from the same litter were pooled to generate one control and one mutant sample, respectively, and resuspended in 400 µl dissociation buffer containing 5% BSA for blocking and incubated for 30 minutes on ice on a horizontal shaker. Alexa Fluor 488 labelled anti-RanBP2 (D-4) antibody (Santa Cruz, sc-74518, RRID:AB_2176784) was added to each sample in a 1:100 final dilution. Samples were incubated for 30 minutes on ice. Wash buffer was added and samples were centrifuged at 850g at 4°C for at least 15 minutes until a solid pellet appeared. The supernatant was decanted and nuclei were resuspended in 180 µl wash buffer. 200 µl 1% BSA in PBS were added and samples were passed through a 40 µm cell strainer before sorting.

Purkinje cell nuclei were isolated by fluorescence-activated nuclei sorting on a BD FACS Aria III cell sorter (BD Biosciences, Heidelberg, Germany) using a 70 µm nozzle and a 1.5 neutral density filter on forward scatter. Single nuclei were gated by side scatter / DAPI signals, and RanBP-positive Purkinje cell nuclei were sorted in a 4-way purity mode. Data were analysed with FACSDiva 9.0.1 software (BD Biosciences, RRID:SCR_001456). 3000-50000 nuclei were sorted into 100 µl TriReagent (Invitrogen, AM8738). Nuclei were stored at -20°C.

#### RNA isolation

RNA was isolated by using the Total RNA Zol Out-D kit by A&A Biotechnology (043-100) and eluted in 25 µl nuclease-free water.

#### Quantitative reverse transcription polymerase chain reaction (RT-qPCR)

To synthesize complementary DNA (cDNA), 12 µl RNA were mixed with 1 µl Random Hexamer Primers (Thermo Scientific SO142) and incubated for 5 minutes at 65°C. 4x reaction buffer, 10 mM dNTPs, 20 U Ribolock (Thermo Scientific EO0381), and 1 µl Revert Ad H Minus M-MuLV Transcriptase (Thermo Scientific EP0451) were added. Samples were incubated at room temperature for 10 minutes, at 42°C for 60 minutes, and reverse transcriptase was inactivated at 70°C for 10 minutes.

For qPCR, cDNA was diluted 1:3 in nuclease free water. Three technical replicates per sample were loaded into a 96 well-plate, with single wells containing the following components: 0.5 µl forward and reverse primers (10 µM), 12.5 µl Power Up SYBR green master mix (Applied Biosystems A25743), 7.5 µl nuclease free water, and 4 µl cDNA template. The qPCR was performed in a StepOne thermal cycler under the following conditions: 10 min 95°C followed by 40 cycles of 95°C for 15 sec and 59°C for 1 min.
The following table displays the primers used in this study:

| **Primer name** | **Sequence (5‘ → 3‘)** |
|---|---|
| *Calbindin* fwd | GACGCTGACGGAAGTGGTTA |
| *Calbindin* rev | GGGTAAGACGTGAGCCAACT |
| *Gap43* fwd | ACCATGCTGTGCTGTATGAGAA |
| *Gap43* rev | CTGAATTTTGGTCGCAGCCT |
| *Ephexin* fwd | GAGGTCTGAACGTGAAGGCA |
| *Ephexin* rev | TGTTCTGTGCGGCTCATCTT |
| *Wasl* fwd | CCCCGCAAGAGAACGAGTC |
| *Wasl* rev | CACCACTGCACTTCTTTGCC |
| *Smarcb1* fwd | ATGGCGTTGAGCAAGACC |
| *Smarcb1* rev | CATACGCAGGTAGTTTCCCAC |
| *Tbp* fwd | GGGAGAATCATGGACCAGAA |
| *Tbp* rev | TTGCTGCTGCTGTCTTTGTT |
| *Gapdh*_fwd | ACTTTGGCATTGTGGAAGGG |
| *Gapdh*_rev | GTAGGAACACGGAAGGCCAT |

Data were analysed using the Step One software (RRID:SCR_014281), Microscope Excel 2016, and GraphPad Prism 9 (RRID:SCR_002798). The ΔΔCt method was used to determine the relative gene expression (RQ):

$$\Delta Ct = Ct\ (gene\ of\ interest) - mean\ Ct\ (reference\ gene\ Gapdh + reference\ gene\ Tbp)$$

$$\Delta\Delta Ct = \Delta Ct\ (experimental\ well) - \Delta Ct\ (control\ group)$$

$$RQ = 2^{-\Delta\Delta Ct}$$

*Gapdh* and *Tbp* served as reference genes, non-Purkinje cells as control group. For statistical analysis, ΔCt-values were used.

#### Behaviour and cognition experiments

The behavioural tests were performed in the following order: open field test, elevated zero test, rotarod, gait analysis, beam walking test, visual cliff test, Morris water maze test. The sequence of behavioural tests was designed to minimise potential confounders related to stress, fatigue, or training effects, and to reduce interference between tasks that assess overlapping behavioural domains. Tests were arranged from less physically demanding and less stressful to more challenging. This order therefore ensured that anxiety and baseline motor activity were measured before motor learning tasks, that simple motor assessments preceded more complex ones, and that cognitively demanding and physically stressful tasks were scheduled at the end of the test battery.

##### Rotarod

The rotarod test was used to investigate the gross motor skills of mice [15]. The rotarod apparatus (TSE Systems) was modified to prevent the mice from changing directions during the test. First, the three open sides of the apparatus were covered to prevent escape and distraction. Furthermore, the width between the spacer discs was narrowed to 3.5 cm and 4 cm for female and male mice, respectively. During the training phase, a mouse was placed on a non-rotating rod with a diameter of 5 cm at a height of 20 cm above the ground for 60 sec (habituation). The mouse was allowed to rest for 10 min in the cage and was placed thereafter twice on the rod rotating at low speed (4 revolutions per minute (rpm)). Following a 30 min break in the cage, the actual test was performed. The mouse was placed on the rod three times for a maximum of 300 sec each time, whereby the speed of the rod rotation was increased linearly every 30 sec from 4 to 40 rpm. The experiment was terminated as soon as the mouse lost its grip and fell off the rod. There was a 15-minutes break between each trial. The following parameters were determined as a measure of motor coordination skills: speed at fall and latency at fall (the time that the animal was able to stay on the rod).

**Gait analysis**

In the gait analysis, paw prints of mice were obtained on paper, and the stride length normalized to body length was determined. The test setup consisted of a tunnel with transparent plastic walls (inner diameter of tunnel: 4.5 cm wide, 40 cm long, 15 cm high), which opened into a box with opaque walls containing food (inner diameter: 10 cm wide, 12 cm long, 9.5 cm high). The floor of the transparent tunnel was lined with paper. The forelimbs and the hindlimbs of a mouse were painted with a non-toxic, water-based paint of two different colours, and the animal was placed in the entrance of the transparent tunnel. Mice prefer dark rooms to light rooms and, therefore, run through the light tunnel into the dark one. The paper with the footprints of the forelimbs and the hindlimbs was used to measure the stride length normalized to the body length.

**Beam walking test**

The beam walking test examines fine motor skills and balance [20]. The animals traversed a 100 cm long elevated narrow beam to reach an escape platform. The level of difficulty was determined by the type of beams used in the test: square beams with a thickness of 6 mm and 12 mm, and round beams with a diameter of 10 mm and 18 mm. The respective beam was fixed 50 cm above the ground. The start platform was brightly illuminated by placing two 60-watt desk lamps, which should trigger a strong flight reflex and cause the animal to move away from the start platform. Two hammocks were placed underneath the stretched beam to pick up a falling mouse during the test and to prevent injuries. A small house with food was present on the target platform. On the day before the first measurement, the mice were trained by placing them on the start platform, which was connected to the target platform via a 25 mm square beam. The trial was repeated at least three times, with 10-minute breaks between each training session. A further repetition was carried out if the animal did not reach the target platform easily. On the next day, the actual test was performed. The crossing of a beam was carried out twice in succession for each type of beam, with a 10-minute break between each test, during which animals were placed in their cage. On a test day, only square or only round beams were used. The following parameters were measured in this test: the time taken to cross the beam, the number of foot slips, and the number of head dips.

**Morris water maze test**

The Morris water maze was used to assess spatial learning and memory [21, 29]. The experiment timeline is shown in Fig. 6h. A water pool, that was filled with turbid water at 22 ± 1 °C, contained an escape platform located below the water surface. External visual cues (patterns, shapes) served as orientation points to locate the platform. The animals were tracked using a camera and video tracking software EthoVision XT (Noldus Information Technology, RRID:SCR_000441). Firstly, the animals were gently accustomed to the test procedure. To this end, they were placed on the platform that was above the turbid water surface, which they usually left themselves after a few seconds and entered the water. Animals were allowed to swim for 15 seconds and to return to the platform. This familiarization phase was carried out three times for each animal. During the initial acquisition, the position of an invisible platform remained the same, and each mouse was placed inside the water pool four times a day from a different starting position. A trial was considered complete as soon as the animal found the platform or after a maximum of 120 s. The animals were trained for 5 days with 4 trials per day (acquisition). After each trial, the animals were allowed to rest for 15-30 minutes under an infrared light in their cage. After the acquisition block, a probe trial I was performed on day 6. To test the reference memory during probe trial I, the escape platform was removed from the pool. The mice were placed opposite of the former platform position and tested for 120 s. Subsequently, a so-called ‘reversal learning’ was carried out, in which the escape platform was moved to the quadrant opposite of the original position. Each mouse was subjected to four reversal trials per day, always starting from different positions, on 5 consecutive days. On day 6, the retention of spatial information was tested again using a probe trial II. The mice were placed opposite of the platform position during the reversal training phase and tested for 120 s. During initial acquisition and reversal training, the time taken to reach the platform was measured on days 1-5. For probe trials I and II, the following parameters were determined: the time spent in the target quadrant, the number of crossings over the platform site, the total distance covered and the mean velocity. The swim paths were manually analysed, and a preferential exploration strategy in each test was identified in accordance with Graziano et al. [12], and Brody and Holtzman [6]. The exploration strategies were classified into thigmotaxis, non-spatial strategies (circling, random searching) and spatial strategies (self-orienting, approaching target, and direct finding) (s. Supplementary Fig. 2).

**Open field test** [13, 24]

Animals were placed near the wall-side of a 50 x 50 cm box with high side walls (approx. 40 cm high) made of non-porous material, and their movements were recorded for a period of 10 min. The recorded video file was analysed using video tracking software EthoVision XT (Noldus Information Technology, RRID:SCR_000441). The mean velocity, the total distance covered during the experiment, and the time spent in central area of the box (25 × 25 cm imaginary square) were measured.

**Elevated zero maze test** [26, 28]

Animals were placed on a ring-shaped platform mounted 50 cm above the floor (inner diameter 40 cm, outer diameter 50 cm) after a 30-minute familiarization phase. This platform was divided into four equally long areas: two opposite open sectors ("arms") without borders and two closed sectors ("arms") surrounded by approximately 12 cm high borders. At the beginning of the experiment, mice were placed in the middle of a closed sector. Their movements were recorded for 30 min and analysed with the video tracking software EthoVision XT (Noldus Information Technology, RRID:SCR_000441). The mean velocity, the total covered distance, the cumulative time spent in open sector, the number of head dips, and the number of animals falling off the apparatus were quantified.

**Visual cliff test**

Visual acuity and depth perception were tested using a visual cliff test [9]. The test setup consisted of a transparent plexiglass chamber measuring 60 x 60 x 19 cm, which was open at the top. A checkerboard-like patterned tissue was placed under the "safe" side of the chamber as well as 60 cm below the "deep" side of the chamber to imitate the cliff drop-off. A bar of 3 cm height and 1 cm width was placed in the centre of the chamber, i.e. at the boundary between the shallow and the feigned deep side. At least one day before the test, vibrissae of mice were trimmed to eliminate the influence of tactile information on choosing a side. During testing, mice were placed in the centre of the elevated bar and allowed to choose between the two sides for a total acquisition time of 5 min and five trials per mouse. The behaviour of mice was recorded by a video camera and analysed with the video tracking software EthoVision XT (Noldus Information Technology, RRID:SCR_000441). The incidence of choosing the "safe" side and the latency to descend from the platform were recorded.

#### Questionnaires and general cognitive function in children with *SMARCB1* mutation

##### Study design and participants

Four children (two boys, two girls) with a confirmed *SMARCB1* pathogenic variant were included. As reference for molecular testing using standard procedures, the *SMARCB1* transcript with accession number NM_003073.3 was used. The mean age of the children was 10.29 years (range = 5.17–13.25). Two individuals (individual B and C) had been previously reported in van der Sluijs et al. (2024) [27], which described their clinical features and MRI findings (patients 41 and 42). The cognitive and behavioural data reported here are novel and have not been presented before.

##### Procedure

Participants were invited for recruitment and assessment by clinicians affiliated with the Leiden University Medical Centre. All evaluations were conducted in the Netherlands. Only children from Dutch-speaking families were included. Children with severe visual or hearing impairments would have been excluded, but this did not apply to any participant. Background information was collected for each child.

##### Behavioural functioning

Behavioural functioning was assessed with the Child Behaviour Checklist (CBCL) [1]. The CBCL is a standardized parent-report questionnaire that evaluates behavioural and emotional problems in children aged 1.5–18 years over the past six months. Chronological age-appropriate versions were used (CBCL 1.5–5 or CBCL 6–18). The questionnaire contains 100 items rated on a 3-point scale (0 = not true, 1 = somewhat/sometimes true, 2 = very true/often true). Higher scores reflect more problems.

Raw scores yield empirically derived syndrome scales as well as DSM-oriented scales. For this study, the syndrome scales were used, as they provide a broad view of behavioural and emotional problems. Scales include: (a) Anxious/Depressed, (b) Withdrawn/Depressed, (c) Somatic Complaints, (d) Social Problems, (e) Thought Problems, (f) Attention Problems, and (g) Rule-Breaking and Aggressive Behaviour. Standardized T-scores were used to interpret results. T-scores above 70 were classified as clinical (++), scores between 65–70 as borderline

(+), and scores below 65 as non-clinical (-). In cases where data were not available, results were coded as N/A.

**Executive behavioural regulation**

Executive behavioural regulation was measured with the Behaviour Rating Inventory of Executive Function, Second Edition (BRIEF-2) [10]. The BRIEF-2 is a parent-report questionnaire designed to evaluate executive functioning behaviours in children aged 5–18 years. It contains 63 items rated on a 3-point scale (0 = never, 1 = sometimes, 2 = often). Higher scores indicate greater executive dysfunction.

The BRIEF-2 yields nine clinical scales grouped into three indices: Behavioural Regulation, Emotional Regulation, and Cognitive Regulation. These cover inhibition, flexibility, emotional control, working memory, and planning/organization. Standardized T-scores were used, with scores above 65 considered clinically elevated (++), scores between 60–65 classified as borderline (+), and scores below 60 as non-clinical (-). If data were not available, this was indicated as N/A.

**Behavioural and emotional functioning**

Behavioural and emotional difficulties were additionally assessed with the Strengths and Difficulties Questionnaire (SDQ) [11]; The SDQ is a brief screening tool for children aged 4–17 years, completed by parents or teachers. It consists of 25 items rated on a 3-point scale (0 = not true, 1 = somewhat true, 2 = certainly true). Items form five subscales: Emotional Symptoms, Conduct Problems, Hyperactivity/Inattention, Peer Relationship Problems, and Prosocial Behaviour.

Cut-offs were applied according to the scoring guidelines of the SDQ. Scores in the abnormal/clinical range (++) indicated significant difficulties, borderline scores (+) indicated moderate difficulties requiring monitoring or further evaluation, and scores in the normal/non-clinical range (-) reflected typical functioning. If information was not available, this was coded as N/A.

**Cognitive functioning: Developmental quotients**

To evaluate the developmental progress of each child, the Developmental Quotient (DQ) was calculated, following procedures used in previous studies [18],[19]. The DQ was obtained by dividing the child's developmental age by their chronological age and multiplying by 100 (DQ = [DA / CA] × 100; M = 100, SD = 15). DQ is particularly useful in research involving young children or individuals with neurodevelopmental disorders, as it enables early detection and monitoring of developmental delays. Previous research has shown a strong linear relationship between DQ and full-scale IQ, supporting its validity as a measure of general cognitive functioning in children [17]. A score of 100 represents development that is typical for a child's chronological age. Scores below 100 indicate developmental delays, whereas scores above 100 suggest more advanced development.

##### Statistical analyses

Shapiro–Wilk test was performed to assess whether the data were normally distributed. Prior to normality tests, the log transformation was applied to the data sets describing width of the primary dendrite in P7 animals and maximal dendritic length in P10 animals. If the data were normally distributed, unpaired t test with Welch's correction was applied to test whether the datasets are significantly different. If the data were not normally distributed or if the normality test could not be applied to the datasets due to low sample size (branching in neurons of supra-pyramidal blade of hippocampus), Mann-Whitney test was used to test whether the datasets are significantly different. Graphs display means and error bars show standard deviations. n.s. not significant ($p > 0.05$), * $p \leq 0.05$, ** $p \leq 0.01$, *** $p \leq 0.001$.
To test whether there is a difference in zone preference (safe zone vs not-safe-zone (=deep side or no descending) choice) between control and mutant mice in visual cliff test, Chi-square test was applied using an online interactive tool [23]. At that, the numbers of mice choosing a safe or not a safe side in at least one of the five trials were considered in the test.
